# A developmental gene regulatory network for invasive differentiation of the *C. elegans* anchor cell

**DOI:** 10.1101/691337

**Authors:** Taylor N. Medwig-Kinney, Jayson J. Smith, Nicholas J. Palmisano, Sujata Tank, Wan Zhang, David Q. Matus

## Abstract

Cellular invasion is a key part of development, immunity, and disease. Using the *in vivo* model of *C. elegans* anchor cell invasion, we characterize the gene regulatory network that promotes invasive differentiation. The anchor cell is initially specified in a stochastic cell fate decision mediated by Notch signaling. Previous research has identified four conserved transcription factors, *fos-1a* (Fos), *egl-43* (EVI1/MEL), *hlh-2* (E/Daughterless) and *nhr-67* (NR2E1/TLX), that mediate anchor cell specification and/or invasive differentiation. Connections between these transcription factors and the underlying cell biology that they regulate is poorly understood. Here, using genome editing and RNA interference, we examine transcription factor interactions prior to and after anchor cell specification. During invasion we identify that *egl-43*, *hlh-2*, and *nhr-67* function together in a type I coherent feed-forward loop with positive feedback. Conversely, prior to specification, these transcription factors function independent of one another to regulate LIN-12 (Notch) activity. Together, these results demonstrate that, although the same transcription factors can function in fate specification and differentiated cell behavior, a gene regulatory network can be rapidly re-wired to reinforce a post-mitotic, pro-invasive state.

**SUMMARY STATEMENT:** Basement membrane invasion by the *C. elegans* anchor cell is coordinated by a dynamic gene regulatory network encompassing cell cycle dependent and independent sub-circuits.

## INTRODUCTION

Invasion through basement membranes (BM) is a cellular behavior that is integral to embryo placentation, tissue patterning during embryonic development, and immune response to infection and injury (Medwig and Matus, 2017; Rowe and Weiss, 2008). Increased cellular invasiveness is also a hallmark of metastatic cancer (Hanahan and Weinberg, 2011). Previous research has identified several cell-autonomous mechanisms that are highly conserved across different contexts of BM invasion (Kelley et al., 2014; Medwig and Matus, 2017). These include localization of F-actin and upregulation of matrix metalloproteinases (MMPs) to chemically degrade the BM (Hagedorn et al., 2009; Hagedorn et al., 2013; Kelley et al., 2019; Levitan and Greenwald, 1998; Morrissey et al., 2014; Sherwood et al., 2005). There is also growing evidence that cells must undergo cell cycle arrest in order to achieve invasive differentiation (Kohrman and Matus, 2017; Matus et al., 2015). How these tightly coordinated programs are transcriptionally regulated is not well understood.

As many contexts of cellular invasion occur deep within tissue layers and are thus difficult to visualize, we utilize morphogenesis of the *C. elegans* uterine-vulval connection as a genetically tractable and visually amenable model for examining cell invasion *in vivo* (Sherwood and Sternberg, 2003). During the mid-L3 stage, a specialized uterine cell called the anchor cell (AC), invades the underlying BM, connecting the uterus to the vulval epithelium to facilitate egg-laying (Fig. 1A,B) (Sherwood and Sternberg, 2003). The AC itself is specified in a cell fate decision event earlier in development, during the L2 stage, where two initially equipotent cells diverge via stochastic Notch asymmetry, giving rise to the terminally differentiated AC and a proliferative ventral uterine (VU) cell (Fig. 1A,B) (Wilkinson et al., 1994).

**Fig. 1.**
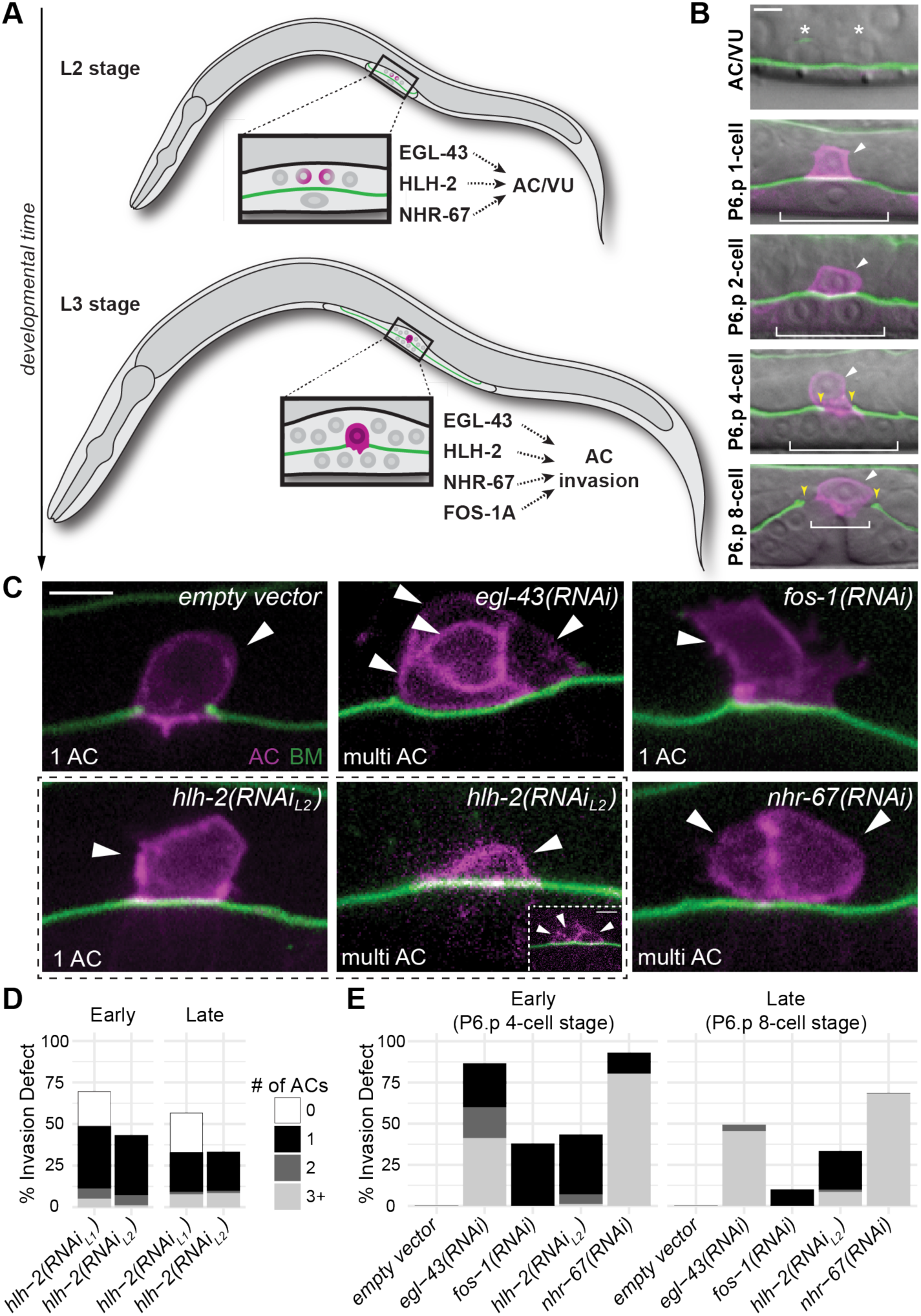
RNAi depletion of pro-invasive TFs leads to defects in AC invasion and proliferation. (A) Schematic depicting the reiterative use of transcription factors in AC specification and invasion. (B) Micrographs depicting AC (magenta, *cdh-3^1.5^*>mCherry::PLCδ^PH^) specification and BM (green, *lam-1*>LAM-1::GFP) invasion over developmental time. (C) Representative empty vector control and TF-RNAi depletion phenotypes, with multiple ACs from additional confocal z-planes shown as insets. (D-E) Stacked bar charts showing the penetrance of invasion defects, binned by number of ACs, following TF-RNAi depletion (p-value < 0.00001, Fisher’s exact test, empty vector compared to each TF RNAi, n ≥ 50 animals examined for each treatment). Due to the penetrance of the zero AC phenotype in early *hlh-2(RNAi)*-treated animals (D), an L2 plating strategy was used to assess post-specification AC invasion phenotypes (E).

Prior research has identified four pro-invasive transcription factors (TFs) that function cell autonomously to regulate AC invasion (Fig. 1A). These include the basic leucine zipper TF *fos-1a* (Fos), the basic helix-loop-helix TF *hlh-2* (E/Daughterless), the nuclear hormone receptor *nhr-67* (NR2E1/Tailless/TLX), and the zinc-finger TF *egl-43* (EVI1/MEL) (Hwang et al., 2007; Matus et al., 2010; Rimann and Hajnal, 2007; Schindler and Sherwood, 2011; Sherwood et al., 2005; Verghese et al., 2011). NHR-67 functions upstream of the cyclin dependent kinase inhibitor, CKI-1 (p21/p27), to induce G1/G0 cell cycle arrest, which is necessary for AC invasion (Matus et al., 2015). Independent of NHR-67 activity, FOS-1A regulates the expression of the long isoform of *egl-43* (EGL-43L) and downstream effectors including multiple MMPs (*zmp-1, -3, and -6*) and a cadherin (*cdh-3*) (Hwang et al., 2007; Kelley et al., 2019; Matus et al., 2010; Rimann and Hajnal, 2007). HLH-2 independently regulates *cdh-3* along with other targets and cytoskeletal polarity (Schindler and Sherwood, 2011). Prior work has also suggested that EGL-43 and HLH-2 may regulate NHR-67 based on conserved HLH-2 binding motifs present in the *nhr-67* promoter as well as *egl-43* and *hlh-2* perturbation experiments using *nhr-67* transcriptional and translational GFP reporters (Bodofsky et al., 2018; Verghese et al., 2011). Interestingly, three of these TFs (*egl-43*, *hlh-2* and *nhr-67*) also play key roles in the specification of the AC and VU cell fates during the L2 stage (Attner et al., 2019; Hwang et al., 2007; Karp and Greenwald, 2004; Sallee et al., 2017; Verghese et al., 2011). How these four TFs function, independently and together, to regulate both specification and invasive activity of the differentiated AC is poorly understood.

Here, using new highly efficient RNA interference (RNAi) vectors (Sturm et al., 2018), we identify novel roles for *egl-43* and *hlh-2*, in maintaining the AC in a cell cycle arrested state. Additionally, we generated GFP knock-in alleles for each pro-invasive TF by inserting GFP in-frame into each genomic locus using CRISPR/Cas9-mediated genome engineering, allowing for quantitative measures of endogenous protein in the context of native chromatin architecture and potential distal enhancer elements that may be misrepresented using traditional transgenic reporters (Dickinson and Goldstein, 2016; Dickinson et al., 2013; Kim et al., 2014). Using these alleles, we report the onset of relative expression of pro-invasive TFs in the lineages leading to the AC and dissect their molecular epistatic interactions. We find that, prior to AC/VU specification, while all four TFs regulate an endogenously tagged *lin-12*::GFP allele, none of the TFs regulate each other. This network appears to be re-wired following specification, as we characterized both a cell cycle independent circuit and a cell cycle dependent feed-forward regulatory circuit with positive feedback critical for maintaining the AC in an invasive, post-mitotic state. These findings provide new insights into how a set of reiteratively used TFs can be re-purposed from regulating cell fate specification to coordinating a differentiated cellular behavior.

## RESULTS

### Improved RNAi penetrance reveals novel phenotypes associated with depletion of pro-invasive TFs

Our initial goal was to determine interactions between the four pro-invasive TFs, *fos-1*, *egl-43*, *hlh-2* and *nhr-67*. To accomplish this, we first generated new RNAi targeting constructs in the improved RNAi vector, T444T, which includes T7 termination sequences to prevent the generation of non-specific RNA fragments from the vector backbone, increasing the efficacy of gene silencing over the original RNAi targeting vector, L4440 (Fig. S1A) (Sturm et al., 2018). As three of four pro-invasive TFs (*egl-43*, *hlh-2* and *nhr-67*) also function during AC specification (Attner et al., 2019; Hwang et al., 2007; Karp and Greenwald, 2004; Sallee et al., 2017; Verghese et al., 2011), we assessed TF depletions in a genetic background where only the AC and neighboring uterine cells are sensitive to RNAi following the AC/VU decision (Matus et al., 2010). The uterine-specific RNAi sensitive strain was generated through tissue-specific restoration of the RDE-1 Piwi/Argonaut protein in an *rde-1* mutant background using the *fos-1a* promoter, which is expressed specifically in the somatic gonad during the late L2/early L3 stage of larval development (Haerty et al., 2008; Hagedorn et al., 2009; Matus et al., 2010; Matus et al., 2015). AC invasion was quantified as the presence or absence of a BM gap, visualized by laminin::GFP, at the P6.p four-cell stage, a developmental window where 100% of wild-type ACs are invaded (Fig. 1C, Table S1). We also assessed invasion several hours later, following vulval morphogenesis, allowing us to distinguish between delays in invasion and complete loss of invasive capacity (Fig. 1E, Fig. S1D, Table S1). Depletion of each TF resulted in invasion defects ranging from moderate penetrance (38%, *fos-1*) to highly penetrant (69-94%, *hlh-2*, *egl*-*43*, and *nhr-67*), when synchronized L1-stage animals were plated on RNAi bacteria and scored at the mid-L3 stage (P6.p 4-cell stage) and early L4 stage (P6.p 8-cell stage) (Fig. 1C-E, Fig. S1D). Consistent with previous work, depletion of *fos-1* resulted in single ACs that failed to breach the BM. Depletion of *nhr-67* resulted in proliferative, non-invasive ACs at a high penetrance. Depletion of *egl-43* phenocopied *nhr-67*, with a highly penetrant defect of multiple, non-invasive ACs. Lastly, depletion of *hlh-2* resulted in pleiotropic phenotypes in the AC, from animals with zero, one, two or even several cases of three or more ACs (Fig. 1D, S1C). In all cases, the penetrance of AC invasion defects was significantly increased using a T444T-based RNAi targeting vector compared to L4440-based vectors (Fig. S1A). Thus, use of the improved RNAi vector revealed novel phenotypes resulting from depletion of *egl-43* and *hlh-2* during invasion, phenocopying depletion of *nhr-67* resulting in multiple non-invasive ACs (Matus et al., 2015).

Due to pleiotropy associated with *hlh-2* depletion, which affects AC specification and invasion (Schindler and Sherwood, 2011), we sought to rule out that the extra *cdh-3*-expressing ACs are due to defects in AC specification. Depletions using the traditional RNAi vector, L4440, in the uterine-specific RNAi-sensitive background is sufficient to bypass AC fate specification defects that occur in genetic backgrounds where all cells are sensitive to RNAi (Matus et al., 2010). Specifically, loss of HLH-2 prior to AC specification disrupts the *lin-12*(Notch)/*lag-2*(Delta)-dependent signaling cascade, resulting in both pre-AC/VU cells adopting a VU fate, resulting in a zero AC phenotype. During AC specification, loss of HLH-2 results in upregulation of pro-AC fate by activating *lag-2*/Delta resulting in both AC/VU cells adopting an AC fate (Karp and Greenwald, 2004; Schindler and Sherwood, 2011). Notably, *hlh-2* depletion using the T444T vector resulted in a significantly greater occurrence of a zero AC phenotype as compared to the L4440 *hlh-2* RNAi clone (Fig. 1D, S1A), suggesting that the enhanced T444T vector may narrow the developmental window between restoration of RDE-1 function, the timing of the AC/VU decision and RNAi sensitivity. Thus, to bypass these potential specification defects due to increased RNAi penetrance for *hlh-2* using the T444T vector, we depleted *hlh-2* and the other TFs by RNAi using an L2-plating strategy (Schindler et al. 2011), growing animals to the time of the L1/L2 molt on OP50 *E. coli* and then transferring them to TF-RNAi plates. L2 platings resulted in a lower overall penetrance of AC invasion defects for all TF-RNAi treatments (Fig. S1B), including *hlh-2* (43% at the P6.p 4-cell stage; Fig. 1D) but all animals possessed an AC. Strikingly, some animals possessed multiple *cdh-3*-expressing ACs early (Fig. 1C), while 10% of animals in the early L4 stage possessed multiple ACs (Fig. 1C-E, Fig. S1C,D). As the number of ACs increased over developmental time (Fig. 1D-E), these results suggest that the multiple ACs observed could be arising through loss of G1/G0-cell cycle arrest and *nhr-67* activity (Matus et al., 2015).

### Characterization of GFP-tagged alleles and RNAi penetrance

Next, we sought to examine the relationship of these four TFs in the AC prior to and during invasion by combining gene knockdown with endogenous GFP-tagged alleles. To accomplish this, we used CRISPR/Cas9 genome editing technology to knock-in a codon-optimized GFP tag into the endogenous locus of each TF (Fig. S2, Tables S4-6) (Dickinson and Goldstein, 2016; Dickinson et al., 2013; Dickinson et al., 2015). First, we examined a minimum of 50 L1/L2-stage animals to determine the onset of expression for each TF in the somatic gonad, which is derived from two founder cells, Z1 and Z4 (Kimble and Hirsh, 1979). The two cells that can stochastically give rise to the AC are the proximal great-granddaughters of Z1 and Z4 (Z1.ppp and Z4.aaa). Examination of L1 and L2 stage somatic gonads revealed that *egl-43*::*GFP::egl-43* turns on first, in Z1 and Z4, and remains on in their descendants through the time of AC/VU specification (Fig. 2A). As reported recently (Attner et al., 2019), *GFP::hlh-2* turns on at the second division of Z1/Z4 cells in Z1.pp and Z4.aa. We also detected onset of *nhr-67::GFP* in these cells in some animals, though it appears that their expression is weaker in Z1.pp/Z4.aa as compared to *GFP::hlh-2*, but becomes robust in the AC/VU cells prior to specification. Finally, we detected weak expression of *GFP::fos-1a* in L1 and L2 Z1/Z4 descendants that increased rapidly in the somatic gonad following AC specification (Fig. 2A). In summary, consistent with previously reported roles for *egl-43*, *hlh-2*, and *nhr-67* in AC/VU specification (Hwang et al., 2007; Karp and Greenwald, 2004; Verghese et al., 2011), we detect the presence of GFP in Z1/Z4 descendants.

**Fig. 2.**
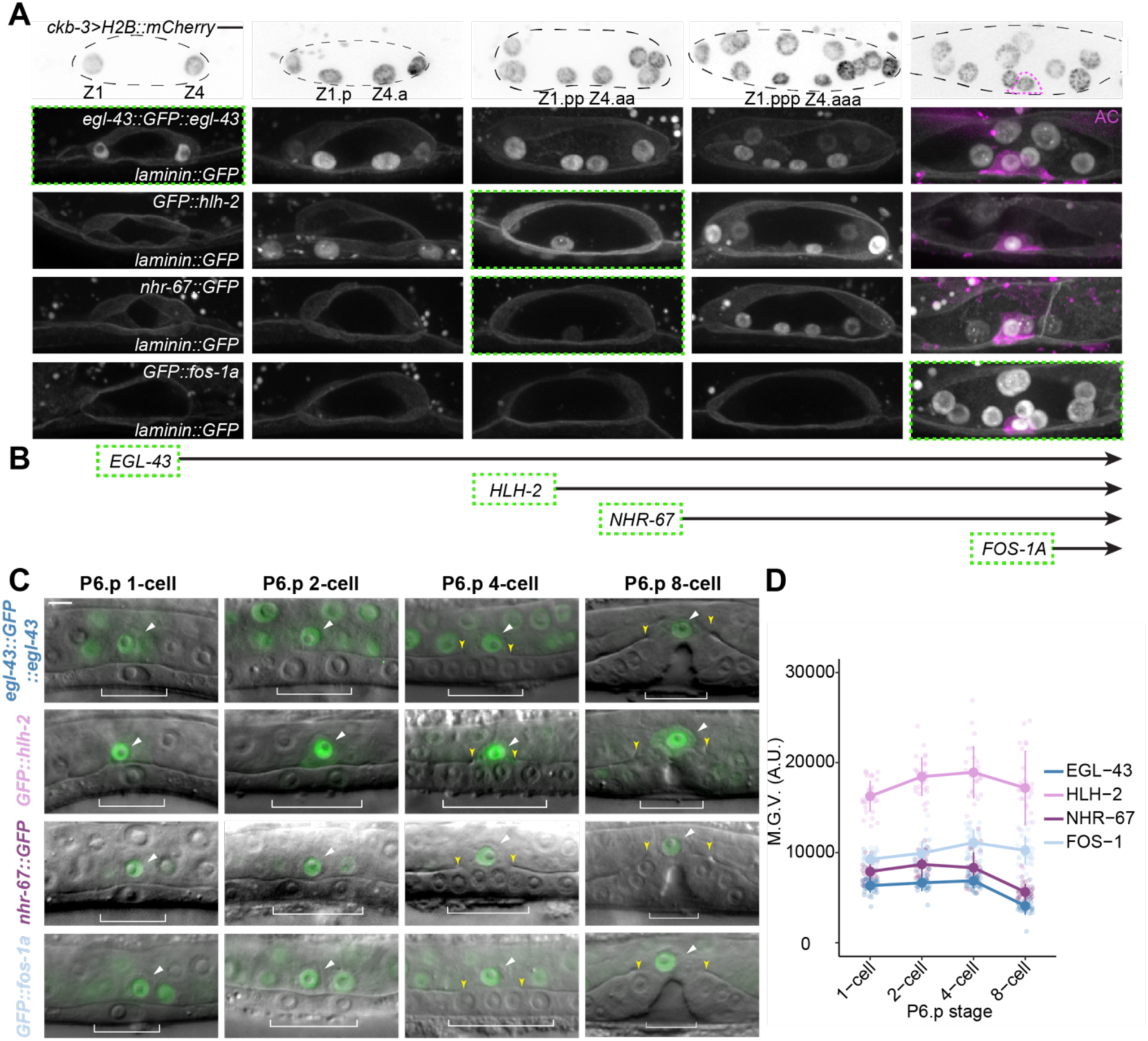
Onset and expression of GFP-tagged alleles of pro-invasive TFs in the AC over developmental time. (A) Maximum intensity projections of GFP-tagged TFs (bottom) in the context of the BM-lined uterus (*laminin*::GFP), in relation to number of somatic cells in the gonad (top, *ckb-3*>H2B::mCherry, inverted). (B) Schematic illustrating the order of onset of pro-invasive TFs over time (n ≥ 25 for each stage). Visualization (C) and quantification (D) of GFP-tagged TF levels in the AC in relation to the division of P6.p. In this and all other figures, circles and error bars denote mean ± s.d. (n ≥ 25 for each stage).

Next, we examined the expression patterns of *TF::GFP* alleles following AC specification. All four TFs showed robust GFP-localization in the AC nucleus prior to and during invasion (Fig. 2B). Additionally, the GFP-tagged strains had similar expression domains in the somatic gonad during the L3 larval stage as previously reported by traditional multi-copy array transgenes (Hwang et al., 2007; Matus et al., 2015; Rimann and Hajnal, 2007; Sherwood et al., 2005; Verghese et al., 2011). After examining expression patterns, we then synchronized TF GFP-tagged strains and collected a developmental time-course quantifying the expression of GFP-tagged protein in the AC during uterine/vulval development (Fig 2B). All GFP-tagged TFs were imaged using uniform acquisition settings on a spinning disk confocal using an EM-CCD camera, allowing us to compare relative expression levels of endogenous GFP-tagged proteins in the AC. Interestingly, all four TFs followed the same general trend in expression levels, with a gradual increase in levels prior to invasion, peaking at or just before the P6.p 4-cell stage when 100% of wild-type animals have generated a gap in the BM. Expression levels then decreased following invasion, during vulval morphogenesis, in the early L4 stage (P6.p 8-cell stage) (Fig. 2C).

As RNAi penetrance can be difficult to measure at single-cell level across a population of animals, we next assessed the efficacy of our newly generated TF-RNAi vectors quantitatively. To accomplish this, we first crossed a BM reporter (*laminin::GFP*) and an AC marker (*cdh-3>mCherry:moeABD*; see Materials and Methods for transgene nomenclature) into each endogenously GFP-tagged TF strain. We then performed a series of RNAi depletion experiments targeting each TF and examining synchronized animals at the P6.p 4-cell stage. As the insertion of GFP-tags into native genomic loci can potentially interfere with gene function, we examined a minimum of 50 animals treated with control (T444T) empty vector RNAi (Fig. 4). All four GFP-tagged strains showed 100% BM breach at the normal time of invasion in control animals (empty vector) (Fig. S3) and the animals all appear superficially wild-type. Additionally, all alleles generated do not appear to affect viability or fertility. Although this does not rule out other more subtle hypomorphic phenotypes, for the purpose of determining TF regulatory relationships during AC invasion, we consider all alleles generated to be equivalent to wild-type AC invasion.

Next, we examined a minimum of 50 animals following targeted TF-RNAi depletions, collecting spinning disk confocal z-stacks using the same acquisition settings as used for the developmental series (Fig. 2B) in all experiments. Following image quantification, our results were consistent with the phenotypes identified in our original uterine-specific RNAi screen (Fig. 1, S3). However, by being able to quantify depletion of endogenous GFP-tagged TF targeted by RNAi, we were more accurately able to correlate phenotype with mRNA depletion. The *nhr-67* and *egl-43* improved RNAi constructs strongly knocked down their GFP-tagged endogenous targets, with depletions averaging 88% and 75%, respectively (Fig. S3E). This strong knockdown of *nhr-67* and *egl-43* was also correlated with highly penetrant AC invasion defects (61% and 100%, respectively). The completely penetrant defect from *egl-43(RNAi)* segregated into 26% of animals exhibiting a single AC that failed to invade and all other cases phenocopying loss of *nhr-67*, with multiple *cdh-3*-expressing non-invasive ACs (Fig. S3E). Depletion of *GFP::fos*-*1a* by RNAi was also highly penetrant, with a mean depletion of 92% of GFP-tagged endogenous protein and 76% of treated animals had a defect in AC invasion (Fig. S3E). Although only loss of *hlh-2* prior to AC specification can affect AC fate directly resulting in zero ACs (Karp and Greenwald, 2004), *nhr-67* and *egl-43* also regulate the AC/VU decision (Hwang et al., 2007; Verghese et al., 2011). Thus, to ensure that the invasion defects we were observing were the result of post-AC specification defects following TF knockdown, we performed L2 platings and scored for invasion. Since L2 platings result in less time (∼12 hours at 25°C) exposed to TF-RNAi, we were not surprised to observe a lower penetrance of AC invasion defects compared to L1 platings targeting the other TFs (Fig. S3F). As all L2 platings phenocopied L1 platings, although at lower penetrance and only depletion of *hlh-2* at the L1 stage directly affects AC fate, we utilized L1 platings for their increased penetrance for *nhr-67*, *egl-43* and *fos-1* for the remainder of our experiments. Together, these results suggest that following strong TF depletion, loss of either *egl-43* or *hlh-2* phenocopy *nhr-67* depletion, generating multiple, non-invasive ACs.

### Identification of a feed-forward loop controlling NHR-67 activity

Next, we examined the relationship between the four pro-invasive TFs during invasion. To accomplish this, we performed a series of RNAi depletion experiments using synchronized L1 or L2 (for *hlh-2*) stage animals and quantified the amount of GFP-tagged endogenous TF in the AC in animals with defects in invasion at the P6.p four-cell stage. While depletion of *fos-1* failed to significantly down-regulate levels of *nhr*-*67::GFP,* depletion of *egl-43* resulted in a strong reduction (65%) of *nhr-67::GFP* in the AC (Fig. 3A,B). Intriguingly, in animals with a single non-invasive AC following *hlh*-*2(RNAi),* we saw only a partial reduction (19%) of *nhr-67::GFP* levels, while in animals with multiple *cdh-3*-expressing cells we saw a significant reduction (49%) in *nhr*-*67::GFP* (Fig. 5A,B). These results suggest that both *egl-43* and *hlh-2* co-regulate *nhr*-*67* during invasion.

**Fig. 3.**
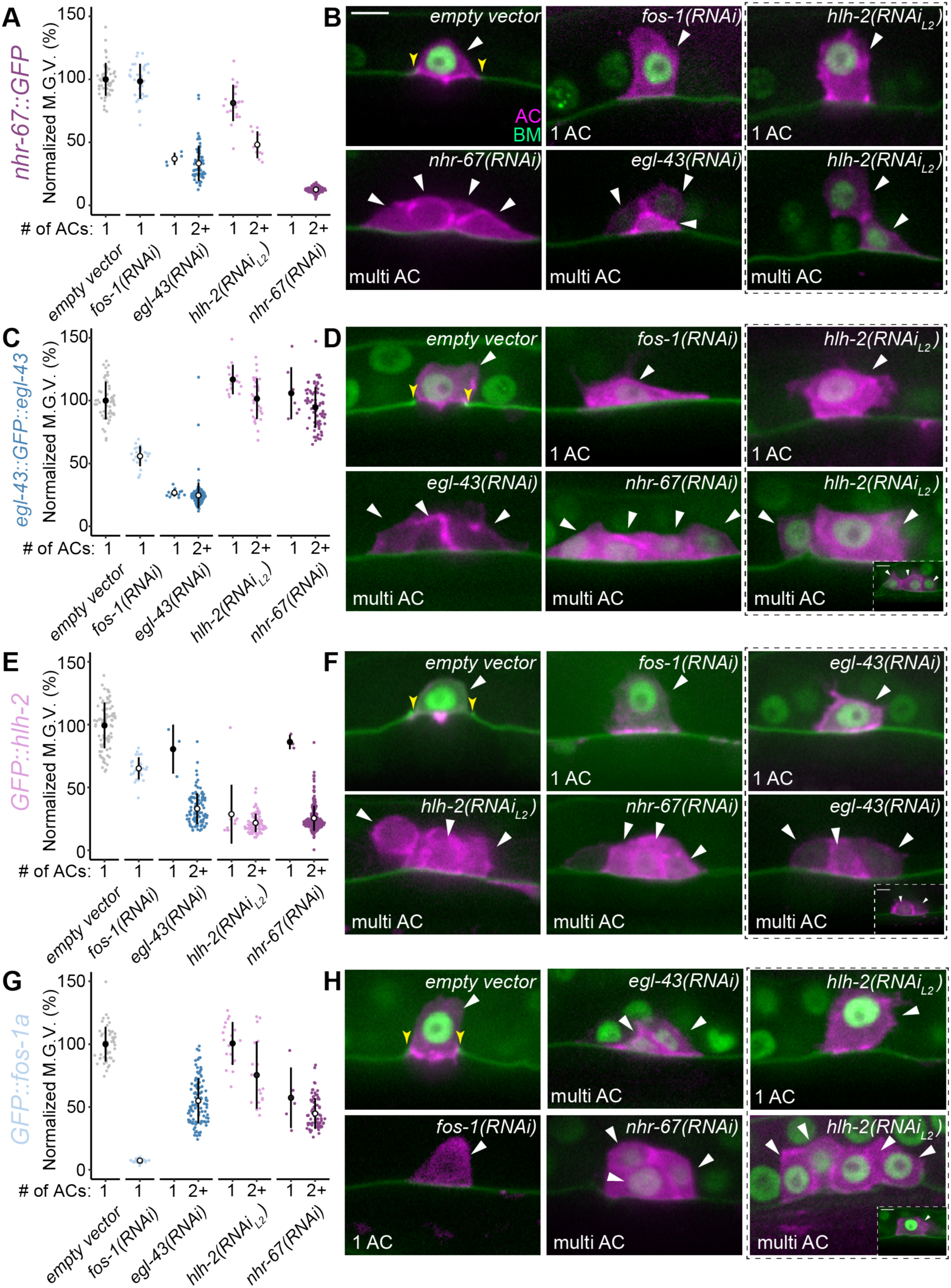
Regulatory interactions among pro-invasive TFs at endogenous loci. (A, C, E, G) Sina plots of GFP-tagged TF levels, defined as the mean gray value of individual AC nuclei, following RNAi perturbation (n ≥ 25 animals per treatment, p < 1×10^-6^, Student’s t-test). (B, D, F, H) Representative phenotypes (single vs. multi AC, as noted bottom left of each image) following TF-RNAi depletion.

**Fig. 4.**
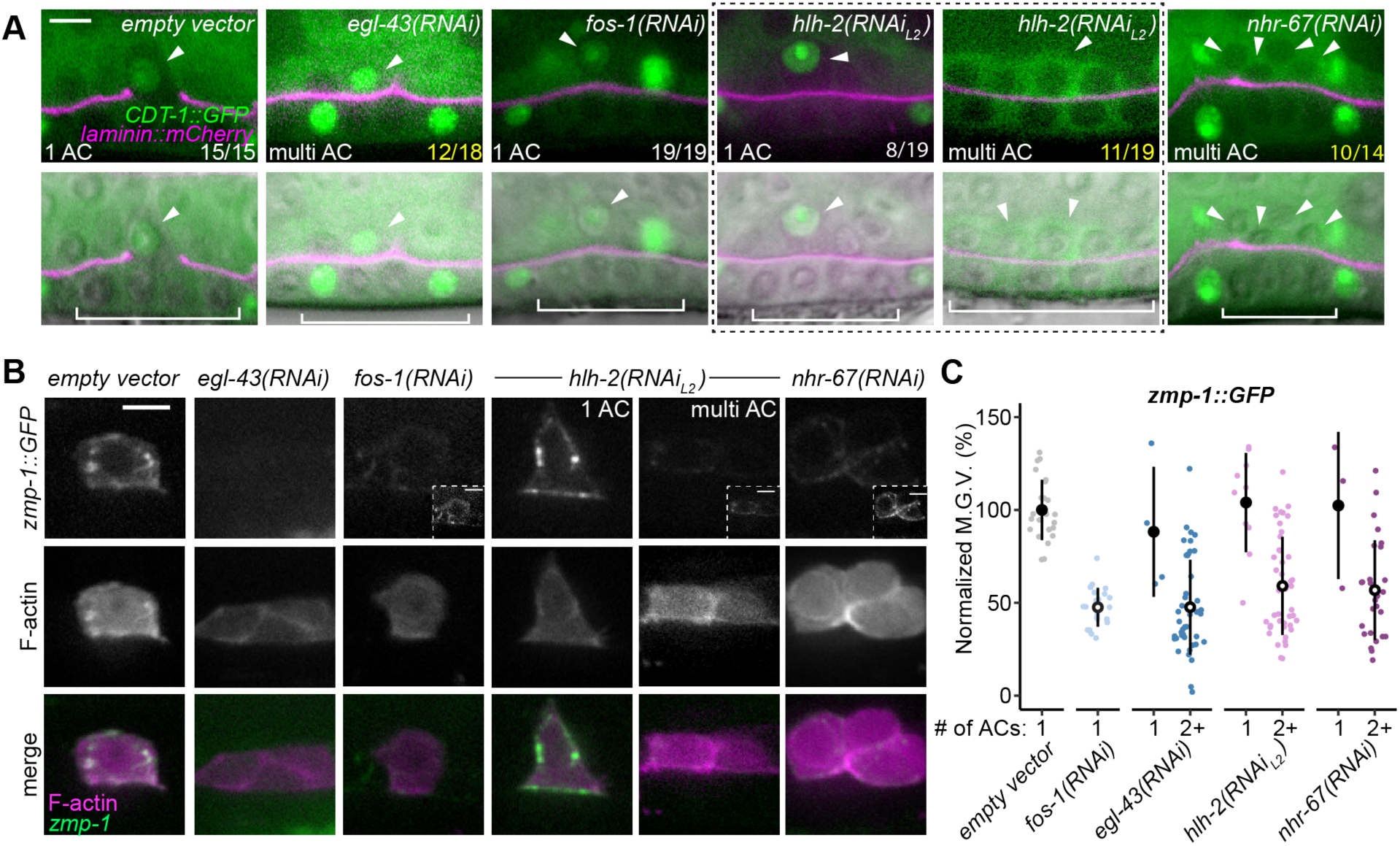
Pro-invasive TFs regulate cell cycle and MMPs. (A) Localization of cell cycle state reporter, *cdt-1*>CDT-1::GFP (green) and BM (*lam-1*>LAM-1::mCherry, magenta) in empty vector control (left) compared to TF-RNAi. White and yellow fractions (bottom right of each micrograph) indicate the number of animals exhibiting the phenotype shown. Visualization (B) and quantification (C) of endogenous MMP (*zmp-1::GFP*) expression in individual ACs following RNAi perturbations (n ≥ 20 animals per treatment, p < 0.01, Student’s t-test).

**Fig. 5.**
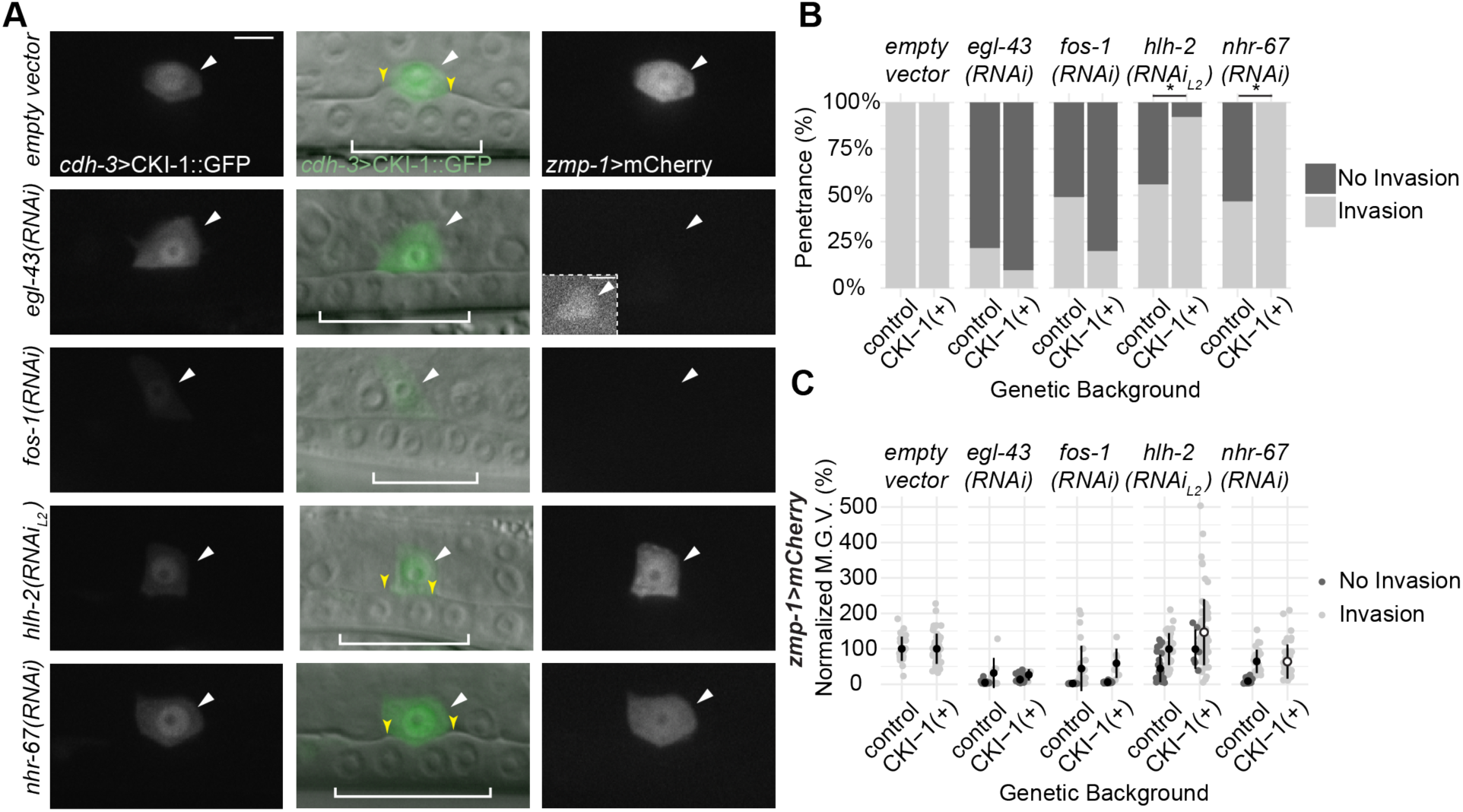
HLH-2 depletion is partially rescued by induced G1/G0 arrest while EGL-43 has a cell cycle-independent pro-invasive role. (A) Micrographs depicting *cdh-*3>CKI-1::GFP (left), DIC overlay (middle), and *zmp-1*>mCherry expression (right) in empty vector control (top) and TF-RNAi depletions. (B) Quantification of AC invasion defects in control and CKI-1(+) animals (n ≥ 27 animals per treatment, p-value < 0.00001, Fisher’s exact test). (C) Quantification of *zmp-1*>mCherry reporter levels in control and CKI-1(+) animals (n ≥ 27 animals per treatment, p-value of < 0.01, Student’s test).

We repeated these TF-RNAi molecular epistasis experiments with the remaining three TF-GFP-tagged strains and quantified depletion of GFP in animals with invasion defects. Consistent with previous studies using transcriptional and translational reporters (Hwang et al., 2007; Rimann and Hajnal, 2007), we found that depletion of *fos-1* regulated levels of *egl-43::GFP::egl-43* (44% depletion) (Fig. 3C,D). No other TF depletion significantly regulated the levels of *egl-43* in our experiments (Fig. 3C,D). *hlh*-2 is predicted to regulate *egl-*43 activity based on the presence of two conserved E-box binding motifs in the *egl-43* promoter (Hwang et al. 2007). However, we did not detect regulation of *egl-43::GFP::egl-43* in either the AC (Fig. 3C,D) or neighboring ventral uterine (VU) cells (Fig. S4). Next, we examined *GFP::hlh-2* levels following TF-RNAi depletions. Intriguingly, we found a similar pattern of regulation based on AC phenotype. Animals with single non-invasive ACs showed partial depletion of *GFP::hlh*-*2* following *fos-1, egl-43,* or *nhr-67* depletion (35%, 20%, and 14%, respectively). However, animals with multiple *cdh-3*-expressing ACs following *egl-43(RNAi)* or *nhr*-*67(RNAi)* showed stronger depletion of *GFP::hlh-2* (66% and 73%, respectively) (Fig. 3E,F). Finally, we examined levels of *GFP::fos-1a* following TF depletions. Depletion of either *nhr-67* or *egl-43* resulted in partial reduction of *GFP::fos-1a* levels in the AC (52% and 49%, respectively) (Fig. 3G,H). Together, these results suggest that *egl-43*, *hlh-2* and *nhr-67* may function together in a feed-forward regulatory loop with positive feedback to maintain the AC in a post-mitotic, pro-invasive state.

### Multiple ACs derive from proliferation

As depletion of *egl-43* and *hlh-2* both regulate levels of *nhr-67* and their depletion results in multiple ACs that fail to invade, we next wanted to assess whether their depletion was functionally phenocopying depletion of *nhr-67*. To confirm that the presence of multiple cells expressing the *cdh-3*-driven AC reporter were due to defects in proliferation, we performed static imaging using a full-length translational *cdt-1>*CDT-1::GFP reporter that indicates cell cycle progression (Fig. 4A). As CDT-1 must be removed from origins of replication during DNA licensing, the transgene localizes to the nucleus during G1/G0 and is largely cytosolic at the onset of S phase (Matus et al., 2014; Matus et al., 2015). As expected, T444T (empty vector) and *fos-1(RNAi)* treated animals consistently exhibited nuclear localization of CDT-1::GFP (Fig. 4A; 100%, T444T, n = 15; 100%, *fos-1(RNAi),* n = 19 animals examined), indicating cell cycle arrest. However, following depletion of *egl-43, hlh-2* and *nhr-67* we identified multiple animals (Fig, 4A) possessing non-invasive ACs lacking nuclear CDT-1::GFP, indicative of cycling ACs (67%, *egl-43(RNAi)*, n = 12/18; 58%, hlh-2(RNAi)L2 n = 11/19; 71%, nhr-67(RNAi), n = 10/14 animals examined). Together, these results strongly support our molecular epistasis data suggesting that *egl-43* and *hlh-2* function upstream of *nhr-67* to maintain the post-mitotic state of the AC during invasion.

After examining the role of pro-invasive TFs in regulating cell cycle progression, we then sought to examine other downstream targets involved in AC invasion. Using an endogenously-tagged GFP reporter for the MMP, *zmp-1* (Kelley et al., 2019), we assessed the ability of ACs to produce MMPs following *TF(RNAi)* depletion. We confirmed previously published findings (Kelley et al., 2019; Sherwood et al., 2005) that *fos-1* depletion results in reduction of *zmp-1* expression (Fig. 4B). Interestingly, in the cases of the other *TF(RNAi)* treatments, a significant reduction of *zmp-1* expression was only observed when multiple *cdh-3*-expressing ACs were present (Fig. 4B), suggesting that cell cycle progression may downregulate production of MMPs.

### EGL-43 and HLH-2 play roles in cell cycle dependent and independent pro-invasive pathways

We have shown previously that the invasive activity of the AC following depletion of NHR-67 can be completely rescued by restoring the AC to a post-mitotic G1/G0 state through upregulation of the cyclin-dependent kinase inhibitor, CKI-1 (p21/p27) (Matus et al., 2015). As our epistasis experiments revealed that *egl-43* and *hlh-2* positively co-regulate NHR-67 activity, we induced AC-specific expression of CKI-1 using a *cdh*-*3*>CKI-1::GFP integrated array and assessed invasion following RNAi depletions (Fig. 5A,B). As expected, induced expression of CKI-1::GFP in the AC completely rescued *nhr-67(RNAi)* treated animals (Fig. 5A,B; 100% invaded (n = 42) as compared to 45% of control animals lacking induced CKI-1::GFP (n = 62)). Additionally, *cdh-3*>CKI-1::GFP partially rescued *hlh-2* depletion in L2 RNAi-feeding experiments by restoring AC invasion in most animals examined (Fig. 5A,B; 87% invaded (n = 77), as compared to 56% of control animals (n = 59)). Inducing G1/G0 arrest in the AC failed to rescue *fos*-*1(RNAi)* treated animals (Fig. 5A,B; 20% invaded (n = 35), as compared to 51% of control animals (n = 53)). AC-specific expression of CKI-1::GFP blocked AC proliferation following *egl-43(RNAi),* however this induced arrest failed to rescue invasion (Fig. 5A,B; 10% invaded (n = 31), as compared to 19% of control animals (n = 37)). This result suggests that *egl-43* has pro-invasive functions outside of the cell cycle-dependent pathway. In support of a role for *egl-43* outside of cell cycle control, induced CKI-1::GFP restores the expression of a reporter of MMP activity (*zmp-1*>mCherry) in *nhr-67*-depleted animals, but neither *fos-1* nor *egl-*43-depleted animals show restoration of *zmp-1* transcription (Fig. 5C). Together, these results reveal the presence of two integrated sub-circuits necessary for invasive activity, with *egl-43* and *hlh-2* likely having a critical role for both circuits.

We next set out to test our putative network topology. To do this, we generated two new single copy transgenic lines expressing either BFP-tagged codon optimized HLH-2 or the genomic region of NHR-67, under the transcriptional control of a heat shock promoter. Induced expression of HLH-2 failed to rescue AC invasion following depletion of *egl-43*, *fos-1*, or *nhr-67*, consistent with *hlh-2* functioning upstream of *nhr-67* and the cell cycle independent roles of *egl-43* and *fos-1* in regulating AC invasion (Fig. S5A). Similarly, induced expression of NHR-67 did not rescue invasion in *egl-43(RNAi)* and *fos-1(RNAi)* treated animals (Fig. S5B). Furthermore, induced expression of NHR-67 failed to rescue AC invasion in *hlh-2(RNAi)* treated animals (Fig. S5B). This result, paired with the prevalence of single non-invasive ACs (Fig. 1C) and only partial rescue from induced CKI-1 (Fig. 5A,B) suggest that *hlh-2* regulates invasive targets independent of its role in regulating *nhr-67* and cell cycle arrest.

As our induced TF experiments failed to rescue invasion defects, we examined an *nhr*-*67* hypomorphic allele (*nhr-67(pf88))*, which we have previously shown results in an incomplete penetrance of mitotic, non-invasive ACs (Matus et al., 2015). Notably, *nhr*-*67(RNAi)* using the enhanced T444T vector did not significantly increase the invasion defect of the hypomorphic allele (Fig. S6), supporting previous results using the L4440-based *nhr-67* RNAi vector (Matus et al., 2015). Thus, the *nhr-67(pf88)* allele may represent a putative null phenotype for AC invasion. Similar to our results with *nhr-67(RNAi)*, depletion of *hlh-2* also failed to significantly enhance the *nhr-67(pf88)* AC invasion defect (Fig. S6). Together, these results suggest that *hlh-2* functions upstream of *nhr-67* in maintaining the AC in a post-mitotic, pro-invasive state.

### EGL-43 isoforms function redundantly and in an autoregulative manner during AC invasion

Two functional isoforms of *egl-43* are encoded in the *C. elegans* genome (Fig. S7A). Previous research has suggested that the longer isoform functions downstream of *fos-1a* to modulate MMP expression and other *fos-1* transcriptional targets, including *cdh-3* (Hwang et al., 2007; Rimann and Hajnal, 2007). The shorter isoform has been predicted to either function in Notch/Delta-mediated AC specification (Hwang et al., 2007) and later Notch/Delta-mediated patterning of the ventral uterus (Hwang et al., 2007) or as a potential competitive inhibitor for long isoform binding of downstream targets (Rimann and Hajnal, 2007). Additionally, *fos-1a* and *egl-43* have been shown to function in an incoherent feed-forward loop with negative feedback, with *fos-1a* positively regulating and *egl-43* negatively regulating the levels of *mig-10*/lamellipodin, a key adhesion protein important for stabilizing the attachment of the AC to the BM (Wang et al., 2014). Titrating levels of MIG-10 is critical, as overexpression of MIG-10 results in AC invasion defects (Wang et al., 2014). Although it is readily apparent that levels of EGL-43 are also important, it is unclear from previous studies whether the two isoforms have divergent functions during invasion. Thus, we next decided to explore *egl-43* isoform function.

First, we were generated a knock-in allele of GFP at the *egl-43* N-terminus to specifically tag the long isoform *(GFP::egl-43L*). This allowed us to compare expression patterns of the long isoform to the internally GFP-tagged allele that should dually label both isoforms (Fig. S7A,B). We examined animals during uterine-vulval development and found overlapping expression patterns between both isoforms with strong AC/VU/DU expression in the somatic gonad (Fig. S7B). Next, we generated an *egl-43L*-specific improved RNAi targeting vector in T444T and examined *egl-43L* depletion in the uterine-specific RNAi sensitive genetic background (Hagedorn et al., 2009; Matus et al., 2010). Depletion of *egl-43L* resulted in a penetrant AC invasion defect, as expected, but notably, we detected the presence of animals with both a single non-invasive AC (19%) and animals with multiple ACs (39%) (Fig. 6A,B), similar to depletion of both isoforms (Fig. 1C,E). We then assessed if the long isoform has the same quantitative relationship to the other TFs as observed when targeting both isoforms. Indeed, *egl-43(RNAi)* and *egl-43L(RNAi)* exhibited comparable levels of depletion of *egl-43::GFP::egl-43* (75%, 78%), *GFP::hlh-2* (66%, 60%), and *nhr-67::GFP* (66%, 64%, respectively) (n ≥ 25 animals examined per treatment) (Fig. S7C).

**Fig. 6.**
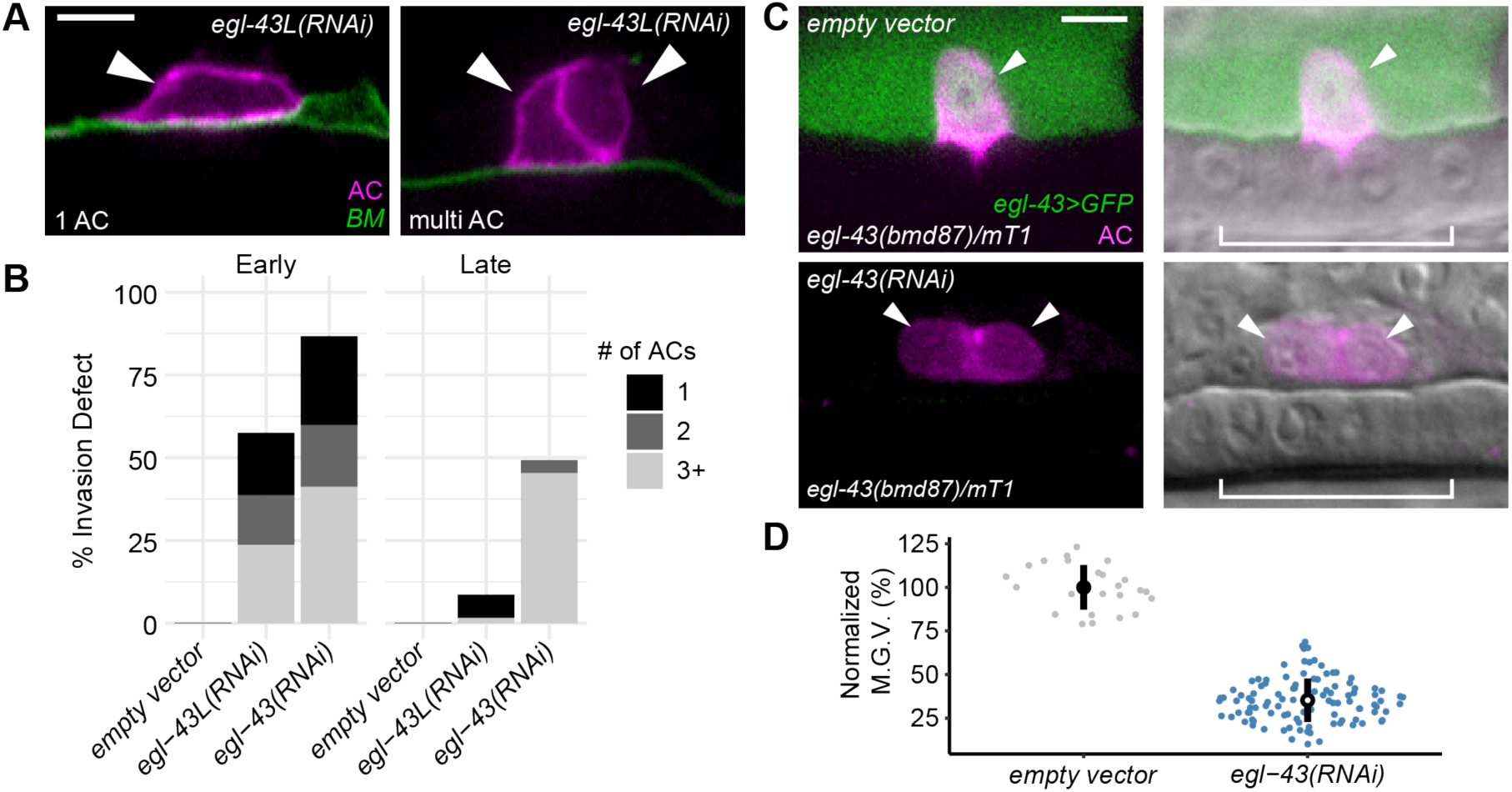
Both isoforms of *egl-43* function redundantly to regulate AC invasion. Visualization (A) and quantification (B) of AC invasion and proliferation defects resulting from RNAi depletion of the long isoform of *egl-43* (*egl-43L*). Visualization (C) and quantification (D) of *egl-43>*GFP expression in individual AC nuclei, assessed using a balanced endogenous *egl-43* transcriptional reporter (*egl-43(bmd87)/mT1*), following *egl-43(RNAi)* treatment compared to control (n ≥ 25 animals per treatment, p < 1×10^-15^, Student’s t-test).

By using the self-excising cassette (SEC) method (Dickinson et al. 2016) to generate GFP knock-in alleles, prior to Cre-Lox recombination, the insertion of GFP at the N-terminus of a locus with a ∼6kb SEC generates a transcriptional reporter at the endogenous locus. Additionally, this ∼6kb DNA cassette also often interferes with the native transcriptional machinery at the locus where it is inserted, generating a strong loss-of-function or putative null allele (Dickinson and Goldstein, 2016). To determine if newly generated alleles exhibit loss-of-function phenotypes and to examine transcriptional output, we next examined the pre-floxed versions of both edited knock-in alleles of *egl-43*, in the presence of an mT1 balancer, as homozygous animals are non-fertile. Similar to the putative null allele, *egl-43(tm1802* (Rimann and Hajnal, 2007), the allele resulting from the SEC insertion of the internal GFP-tag had an extremely low frequency of escapers, and we were unable to obtain L3-stage animals to examine for AC invasion defects. However, similar to our results with RNAi (Fig. 6A,B), an allele resulting from SEC insertion with the long isoform (*egl-43(bmd135)*) displayed a 31% AC invasion defect (n = 11/36), with animals containing either single or multiple non-invading ACs (Fig. S7D,E). Finally, as previous reports based on transgenic transcriptional reporters have suggested that *egl-43* may be autoregulatory (Matus et al., 2010; Rimann and Hajnal, 2007; Wang et al., 2014), we examined the mT1-balanced pre-floxed allele of the internally-tagged *egl-43* targeting both isoforms (*egl*-*43::GFP^SEC^::egl-43*). In support of these previous studies we found strong evidence of autoregulation at the endogenous locus, as *egl-43(RNAi)* reduced the expression of GFP by 65% (n ≥ 25 animals) (Fig. 6C,D). Taken together, these results suggest that both isoforms of *egl-43* function redundantly to regulate multiple transcriptional sub-circuits critical for invasion.

### Activation of the pro-invasive gene regulatory network (GRN) occurs post-specification

Our results here identify a putative feed-forward loop between *egl-43*, *hlh-2* and *nhr-67* to maintain levels of NHR-67, that we have previously shown is critical for keeping the AC in a post-mitotic, pro-invasive state (Matus et al. 2015). As these same TFs also have been shown to function during the AC/VU decision (Hwang et al., 2007; Karp and Greenwald, 2004; Verghese et al., 2011) we examined if the same regulatory relationships were utilized during cell fate specification and differentiation. Intriguingly, our examination of TF onset establishes a hierarchy of *egl-43*, *hlh-2*, and *nhr-67* reaching steady-state levels in the descendants of Z1 and Z4, although all three TFs are robustly expressed in the AC/VU cells (Fig. 2A). To examine TF regulatory relationships prior to specification, we performed L1 TF RNAi platings and examined expression levels of TF-GFP in the AC/VU cells in L2 stage animals. While each TF RNAi robustly depleted its target GFP allele, we failed to detect any molecular epistatic interactions between TFs in the AC/VU cells (Fig. 7A,B). In support of previous research demonstrating a role for these TFs in regulating the AC/VU decision (Hwang et al., 2007; Karp and Greenwald, 2004; Verghese et al., 2011), we detected significant depletion of an endogenously GFP-tagged allele of *lin-12* (Notch) (Fig. 7C,D). Together, these results suggest that despite the re-iterative use of the same set of TFs in specification and invasion, the GRN we characterize here is specific for the post-specification pro-invasive behavior of the AC.

**Fig. 7.**
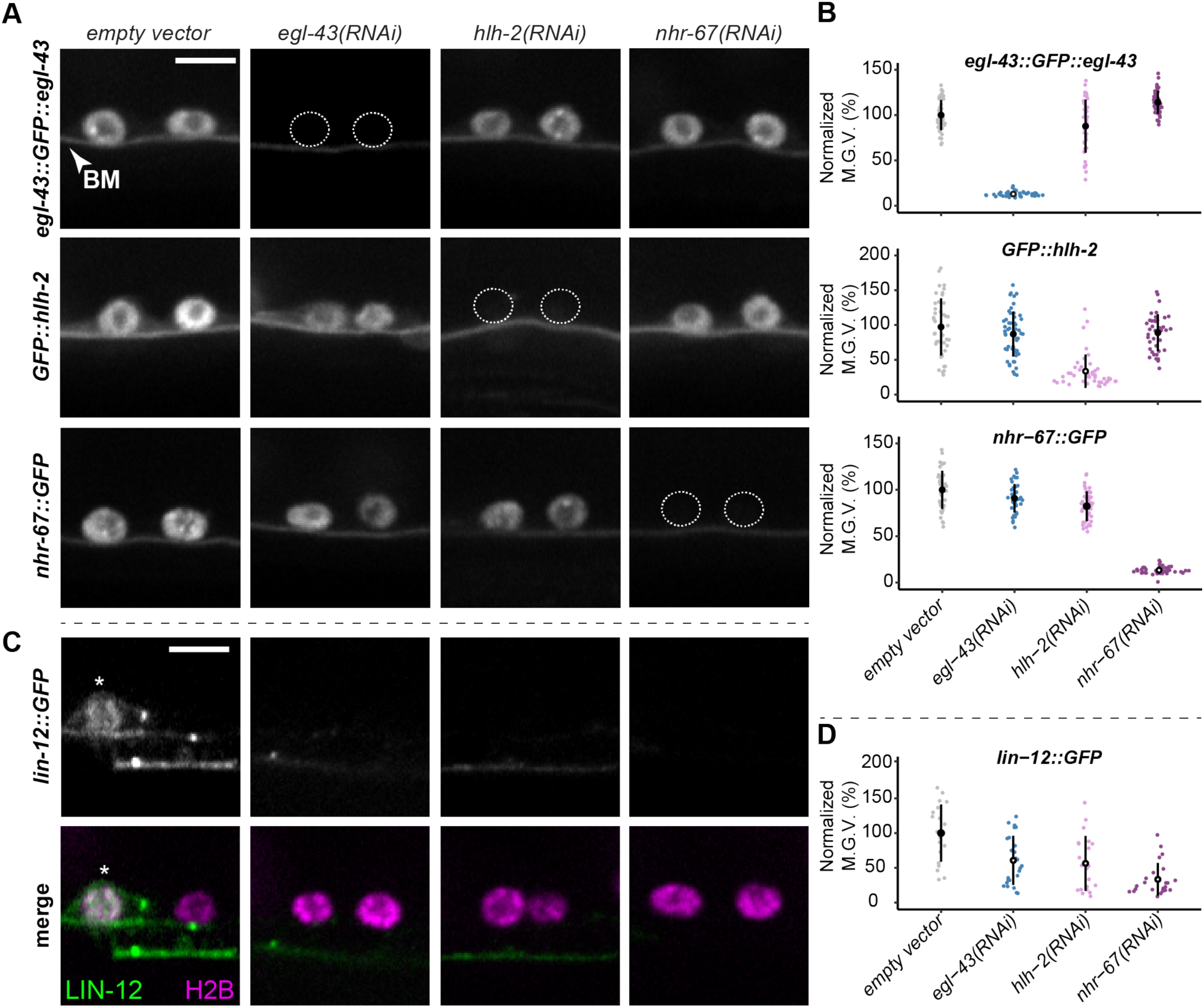
Regulatory relationships do not exist prior to AC specification. Visualization (A) and quantification (B) of GFP-tagged TF levels in the AC/VU precursors (Z1.ppp and Z4.aaa) following RNAi perturbations (n ≥ 20 animals per treatment, p < 1 × 10^-6^, Student’s t-test). Visualization (C) and quantification (D) of GFP-tagged *lin-12* in the AC/VU precursors following RNAi perturbations (n ≥ 20 animals per treatment, p < 0.01, Student’s t-test).

## DISCUSSION

Cellular invasion requires coordination of extrinsic cues from the surrounding microenvironment and orchestration of intrinsic regulatory circuits (Rowe and Weiss, 2008; Sherwood and Plastino, 2018). We focus here on exploring the intrinsic pathways that autonomously regulate *C. elegans* AC invasion. Combining improved RNAi targeting vectors with GFP-tagged alleles enabled us to correlate phenotype to depletion of the endogenous mRNAs and compare regulatory relationships in a quantitative framework. Our experiments reveal new roles for *egl-43* and *hlh-2* in regulating *nhr-67*-mediated cell cycle arrest and suggest that these TFs function in a coherent feed-forward loop with positive feedback. Additionally, we confirm previous work demonstrating that *egl-43* is autoregulatory and demonstrate that it functions in both cell cycle-dependent and independent sub-circuits to orchestrate invasion (Fig. 8). Together, building upon previous work using transgenic approaches (Bodofsky et al., 2018; Hwang et al., 2007; Rimann and Hajnal, 2007; Schindler and Sherwood, 2011; Verghese et al., 2011), our results using endogenously tagged alleles provide the first description of the native regulatory relationships between the four TFs that promote invasive differentiation during *C. elegans* uterine-vulval attachment.

**Fig. 8.**
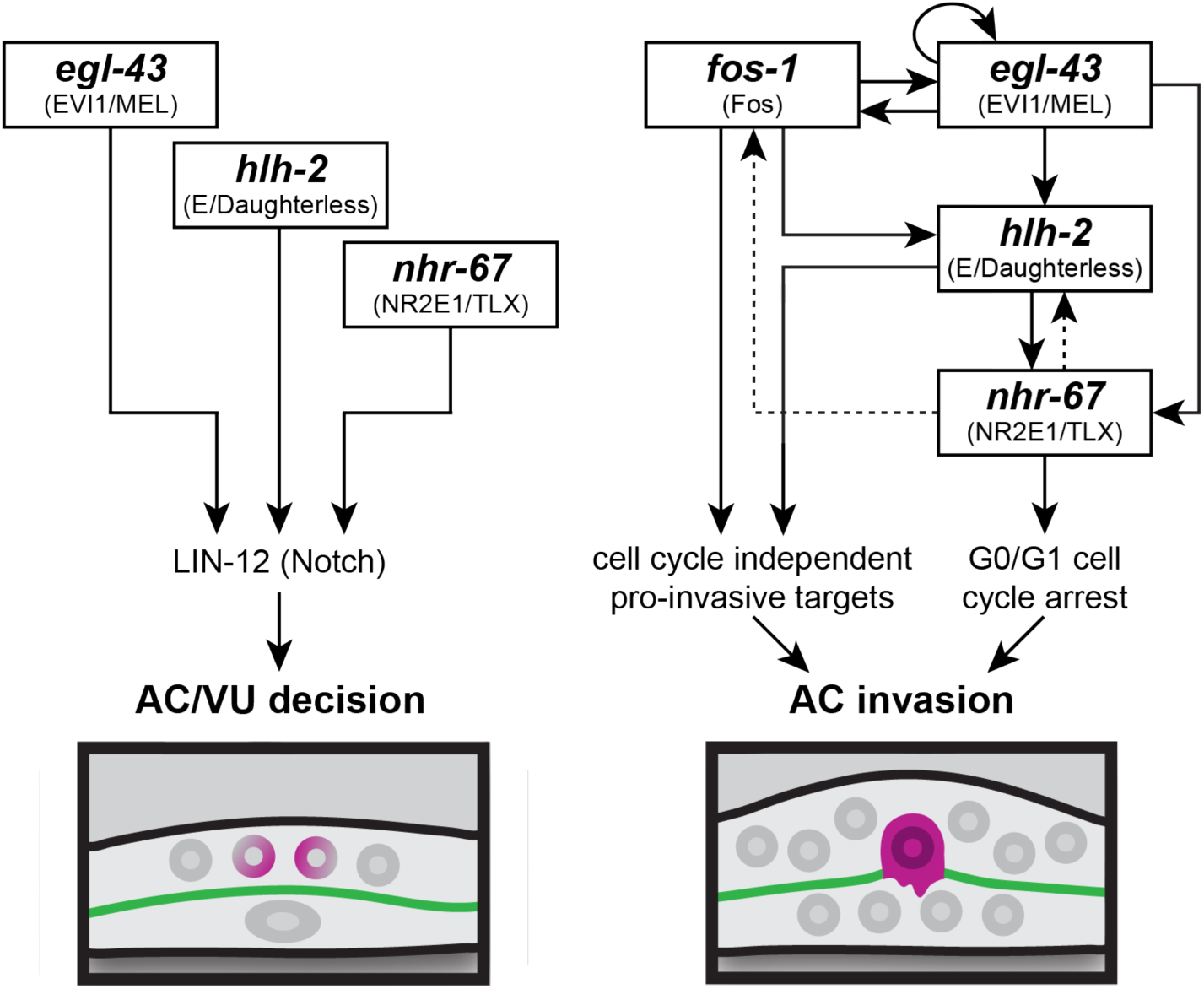
Summary model of the GRNs coordinating AC specification and invasion. The Notch-mediated AC/VU cell fate decision appears to be independently regulated by *egl-43*, *hlh-2*, and *nhr-67* (left). Network inference predicts the GRN regulating AC invasion contains cell cycle independent and dependent sub-circuits, the latter entailing a type I coherent feed-forward loop with positive feedback to maintain the AC in a post-mitotic, invasive state (right). Dotted lines represent predicted feedback.

A key function of *nhr-67* is to maintain the post-mitotic state of the AC (Matus et al., 2015). Here, we demonstrate that *egl-43* and *hlh-2* both regulate *nhr-67*, and that *nhr*-*67*-depletion also reduces HLH-2 levels. Network inference suggests these three TFs function in a classic type I coherent feed-forward loop regulatory circuit with positive feedback (Mangan and Alon, 2003) (Fig. 8). Our model fits with data by others showing that *hlh-2* may directly bind to the *nhr-67* promoter through two canonical E-box motifs found in a 164 bp window in a functional promoter element deleted in several hypomorphic alleles of *nhr-67* (Bodofsky et al., 2018; Matus et al., 2015; Verghese et al., 2011). Further, deletion of these *hlh-2* conserved binding sites results in loss of AC-specific GFP expression in transgenic lines (Bodofsky et al., 2018). While others have also reported positive regulation by EGL-43 on *nhr-67* activity via transgenic reporters (Bodofsky et al., 2018), this interaction, and many of the others TF interactions, may be indirect, as no known binding sites exist for EGL-43 in the *nhr-67* promoter. Alternatively, EGL-43 and NHR-67 have been predicted, based on yeast two-hybrid experiments, to interact at the protein-protein level (Reece-Hoyes et al., 2013). We have also examined the activity of the two previously identified isoforms of *egl-43* (Hwang et al., 2007; Rimann and Hajnal, 2007). As both mutant analyses and RNAi depletion of the long isoform phenocopy depletion of both isoforms, our data suggest that either the long isoform of *egl-43* functions redundantly with the short isoform or that it functions as the dominant isoform at the intersection of multiple regulatory circuits. Of the these two possible hypotheses, the latter is supported by a recent report demonstrating that the long isoform of *egl-43* is a key upstream regulator of the post-mitotic state of the AC as mutation of the transcriptional start methionine of the short isoform of *egl-43* has no observable AC invasion defect and an AC-specific genetic knockout of the long isoform results in mitotic, non-invasive ACs (Deng et al., 2019).

Together, our data and corroborating evidence from the literature support the existence of a coherent feed-forward circuit with positive feedback among *egl-43*, *hlh-2* and *nhr*-*67,* in maintaining the post-mitotic state of the AC (Fig. 8). We have previously shown that *nhr-67* positively regulates transcripts of *cki-1* specifically in the AC (Matus et al., 2015). Thus, to test our putative regulatory circuit, we induced expression of *cki-1* in the AC, which prevents the AC from inappropriately entering the cell cycle. This forced G1/G0 arrest fully rescues *nhr-67* depletion (Matus et al., 2015). Induced *cki-1* partially rescued *hlh-2* depletion but failed to rescue *egl-43* depletion. Heat shock induced expression of *nhr-67* failed to rescue either *hlh-2* or *egl-43*, suggesting that *hlh-2*, like *egl-43*, has a cell cycle-independent function. These results suggest that *egl-43* and *hlh*-*2* function in multiple regulatory circuits. For *egl-43*, this is supported by recent work demonstrating a type I incoherent feed-forward loop between *fos-1a, egl-43L* and the BM adhesion protein MIG-10/lamellipodin (Wang et al., 2014). Finally, our observation that *egl-43* is autoregulatory, also shown previously by *promoter*>GFP fusion experiments (Matus et al., 2015; Rimann and Hajnal, 2007; Wang et al., 2014) supports a model where *egl-43* occupies a key node at the intersection of multiple pro-invasive circuits (Fig. 8).

Feed-forward regulatory loops are likely the most well described network motif occurring in all GRNs (Cordero and Hogeweg, 2006; Davidson, 2010; Mangan and Alon, 2003). From *E. coli* (Milo et al., 2002; Shen-Orr et al., 2002) and yeast (Lee et al., 2002; Milo et al., 2002) to a myriad of examples across the Metazoa (reviewed in Davidson, 2010), feed-forward loops are thought to function as filters for transient inputs (reviewed in Alon, 2007; Hinman, 2016). The addition of positive feedback, generating a coherent feed-forward loop, provides stability to a sub-circuit and is often found in differentiation gene batteries coincident with autoregulation (Davidson, 2010). These network motifs have been described in many developmental contexts, from MyoD-driven vertebrate skeletal muscle differentiation (Penn et al., 2004), the patterning of the Drosophila egg shell via the TF Broad and interactions with EGFR and Dpp signaling (Yakoby et al., 2007), Pax6-dependent regulation of c-Maf and crystallin expression in the mouse embryonic lens (Xie and Cvekl, 2011) and terminal selector neuronal differentiation in *C. elegans* and mammals (reviewed in Hobert, 2008). During *C. elegans* embryonic development, a series of coherent feed-forward loops utilizing SKN-1/MED-1,2 and then a suite of reiteratively used GATA factors (END-1,-3) are required to successfully pattern endomesoderm development (reviewed in Maduro, 2009). Thus, our identification of a coherent feed-forward loop in the *C. elegans* AC maintaining the post-mitotic state, likely evolved to coordinate AC cell cycle exit, both as a by-product of terminal differentiation and a morphogenetic requirement for invasive behavior (Matus et al., 2015). AC invasion is necessary for egg-laying, and defects in the process results in penetrant Protruding vulva/Egg-laying defective (Pvl/Egl) phenotypes, reducing fecundity in individual animals by nearly ten-fold. Thus, redundant control of the sub-circuit regulating differentiation and cell cycle arrest may have been under strong selection, providing an explanation for the regulatory relationships between *egl-43*, *hlh-2* and *nhr-67* we characterize here.

As *egl-43*, *hlh-2* and *nhr-67* reach steady state levels prior to the time of AC/VU specification and individually have been previously shown to regulate different aspects of the stochastic cell fate decision between the AC and VU (Hwang et al., 2007; Karp and Greenwald, 2004; Verghese et al., 2011), we performed RNAi depletion experiments and examined TF::GFP levels in the AC/VU cells during the L2 stage. Surprisingly, we found no evidence of regulation between the TFs at this stage, though as expected, all three TFs regulated levels of endogenously tagged *lin-12::GFP* (Notch). This suggests that the feed forward loop functions specifically in the post-specification AC to maintain the post-mitotic, invasive state (Fig. 8). One plausible explanation for this change in network topology between specification and differentiation is that coordination between chromatin remodeling and TF activity likely changes the genomic landscape between the bipotential AC/VU cells and the post-mitotic, differentiated AC. Additionally, there may be differentially expressed co-factor(s) functioning during specification or invasion that facilitate interactions between TFs. Reiterative use of the same TFs to program different cell behaviors has been examined during neuronal differentiation (Guillemot, 2007) and neural crest specification and migration, as the winged-helix TF FoxD3 was recently shown to regulate multiple independent chromatin-organizing circuits, functioning as a pioneer factor during zebrafish neural crest specification and a transcriptional repressor during later events, including EMT and migration (Lukoseviciute et al., 2018). Whether or not *egl-43*, *nhr-67* or *hlh-2* function as pioneer factors during AC/VU specification is an open question, and one will soon be tractable through FACS isolation and the application of newer next-generation sequencing tools such as ATAC-seq (Assay for Transposase-Accessible Chromatin using sequencing) or CUT&RUN (Cleavage Under Targets and Release Using Nuclease) (Skene and Henikoff, 2017).

We are just beginning to understand connections between regulatory circuits and morphogenetic behaviors (Christiaen et al., 2008; Martik and McClay, 2015; Saunders and McClay, 2014). It is our hope that the ease of genome editing and protein perturbation strategies will facilitate the kind of analyses we have performed here in other metazoan systems. In summary, in this study we characterize the complex relationships between the four pro-invasive TFs that function during AC differentiation to program invasive behavior. We show that although the TFs are reiteratively used in both cell fate specification and in a differentiated cell behavior, invasion, that the network topology is rewired following specification. We identify a classic type I feed forward loop regulating mitotic exit and controlling a switch between proliferative and invasive behavior. Whether or not similar circuit architecture is utilized to regulate invasive and proliferative cell biology in other developmental invasive contexts, including mammalian trophoblast implantation and placentation (Carter et al., 2015; Red-Horse et al., 2004) and epithelial to mesenchymal transition events during gastrulation (Vega et al., 2004) is still poorly understood. Finally, as there appear to be many cancer sub-types that may switch between proliferative and invasive fates (reviewed in Kohrman and Matus, 2017), improving our understanding of the transcriptional network architecture of invasive cells may provide new therapeutic nodes to target in reducing the lethality associated with cancer metastasis.

## MATERIALS AND METHODS

### *C. elegans* strains and culture conditions

Animals were reared under standard conditions and cultured at 25°C, with the exception of temperature-sensitive strains including those containing heat shock inducible transgenes or the *rrf-3(pk1426)* allele (conferring RNAi hypersensitivity), which were maintained at 15°C and 20°C (Brenner, 1974; Simmer et al., 2002). Heat shock inducible transgenes were activated by incubating animals on plates at 32°C for 1 hour following AC specification and again just prior to AC invasion. Animals were synchronized for experiments through alkaline hypochlorite treatment of gravid adults to isolate eggs (Porta-de-la-Riva et al., 2012). In the text and figures, we designate linkage to a promoter through the use of the (>) symbol and fusion of a proteins via a (::) annotation. The following transgenes and alleles were used in this study: *qyIs102*[*fos*-*1>RDE-1;myo-2>YFP*] **LG I** *hlh-2(bmd90*[*hlh-2>LoxP::GFP::HLH-2*]*), qyIs227*[*cdh-3>mCherry::moeABD*]*, bmd121*[*LoxP::hsp>NHR-67::2xBFP*]*, bmd142*[*hsp>HLH*-*2::2xBFP*]; **LG II** *egl-43(bmd87*[*egl-43>SEC::GFP::EGL-43*]*), egl-43(bmd88*[*egl-43>LoxP::GFP::EGL-43*]*), egl-43(bmd136*[*egl-43L>LoxP::GFP::EGL-43*]*) rrf-3(pk1426); qyIs17* [*zmp-1>mCherry*] **LG III** *unc-119(ed4),* **LG IV** *nhr-67(syb509*[*nhr-67>NHR*-*67::GFP*]*), qyIs10*[*lam-1>LAM-1::GFP*] **LG V** *fos-1(bmd138*[*fos-1>LoxP::GFP::FOS-1*]*), qyIs225*[*cdh-3>mCherry::moeABD*]*, rde-1(ne219), qyIs24*[*cdh-3^1.5^>mCherry::PLCδPH*]*, qyIs266*[*cdh-3>CKI-1::GFP*] **LG X** *qyIs7*[*lam-1>LAM-1::GFP*]. See Table S3 for additional details of strains used and generated in this study.

### Molecular biology and microinjection

Transcription factors were tagged at their endogenous loci using CRISPR/Cas9 genome editing via microinjection into the hermaphrodite gonad (Dickinson and Goldstein, 2016; Dickinson et al., 2013). Repair templates were generated as synthetic DNAs from either Integrated DNA Technologies (IDT) as gBlocks or Twist Biosciences as DNA fragments and cloned into *ccdB* compatible sites in pDD282 by New England Biolabs Gibson assembly (Dickinson et al., 2015). Homology arms ranged from 690-1200 bp (see Tables S4-6 for additional details). sgRNAs were constructed by EcoRV and NheI digestion of the plasmid pDD122. A 230 bp amplicon was generated replacing the sgRNA targeting sequence from pDD122 with a new sgRNA and NEB Gibson assembly was used to generate new sgRNA plasmids (see Tables S4-6 for additional details). Hermaphrodite adults were co-injected with guide plasmid (50 ng/μL), repair plasmid (50 ng/μL), and an extrachromosomal array marker (pCFJ90, 2.5 ng/μL), and incubated at 25 °C for several days before carrying out screening and floxing protocols associated with the SEC system (Dickinson et al., 2015).

### RNA interference

An RNAi library of the pro-invasive TFs was constructed by cloning 950-1000 bp of synthetic DNA (663 bp for the *egl-43* long-specific isoform) based on cDNA sequences available on WormBase (www.wormbase.org) into the highly efficient T444T RNAi vector (Grove et al., 2018; Sturm et al., 2018). Synthetic DNAs were generated by IDT as gBlocks or Twist Biosciences as DNA fragments and cloned into restriction digested T444T using NEB Gibson Assembly (see Tables S4-5,7 for additional details). For most experiments, synchronized L1 stage animals were directly exposed to RNAi through feeding with bacteria expressing dsRNA (Conte Jr. et al., 2015). Due to the fact that early *hlh-2* RNAi treatment perturbs AC specification, animals were initially placed on empty vector RNAi plates and then transferred to *hlh-2* RNAi plates following AC specification, approximately 12 hours later at 25°C or 24 hours later at 15°C (Schindler and Sherwood, 2011).

### Live cell imaging

All micrographs included in this manuscript were collected on a Hamamatsu Orca EM-CCD camera mounted on an upright Zeiss AxioImager A2 with a Borealis-modified CSU10 Yokagawa spinning disk scan head using 488 nm and 561 nm Vortran lasers in a VersaLase merge and a Plan-Apochromat 100x/1.4 (NA) Oil DIC objective. MetaMorph software (Molecular Devices) was used for microscopy automation. Several experiments were scored using epifluorescence visualized on a Zeiss Axiocam MRM camera, also mounted on an upright Zeiss AxioImager A2 and a Plan-Apochromat 100x/1.4 (NA) Oil DIC objective. Animals were mounted into a drop of M9 on a 5% Noble agar pad containing approximately 10 mM sodium azide anesthetic and topped with a coverslip.

### Scoring of AC invasion

AC invasion was scored at the P6.p 4-cell stage, when 100% of wild-type animals exhibit a breached BM (Sherwood and Sternberg, 2003). Presence of green fluorescence under the AC, in strains with the laminin::GFP transgene, or presence of a phase dense line, in strains without the transgene, was used to assess invasion. Wild-type invasion is defined as a breach as wide as the basolateral surface of the AC, whereas partial invasion indicates the presence of a breach smaller than the footprint of the AC (Sherwood and Sternberg, 2003). Raw scoring data is available in Tables S1-2.

### Image quantification and statistical analyses

Images were processed using FIJI/ImageJ (v.1.52q) (Schindelin et al., 2012). Expression levels of TFs were quantified by measuring the mean gray value of AC nuclei, defined as somatic gonad cells strongly expressing the *cdh*-*3*>mCherry::moeABD transgene, subtracted by the mean gray value of a background region of equal area to account for EM-CCD camera noise, which we use as a proxy for background fluorescence, as measurements of the mean grey value of unlabeled ACs approximately correspond to camera noise (means = 2514.2 and 2243.5, respectively). Levels of *zmp-1*>mCherry were quantified in control and *cdh-3>*CKI-1::GFP animals by measuring the mean gray value of the entire AC, selected either by a hand drawn region of interest or using the threshold and wand tools in FIJI/ImageJ. For characterization of downstream targets (Fig. 4) and molecular epistasis experiments (Fig. 3, S3 & S7) at the L3 stage, only animals TF-RNAi exhibiting defects in invasion were included in the analysis. Data was normalized to negative control (empty vector) values for the plots in Fig. 3-7, S5-6, & S7, and to both negative control and positive control values for determining the interaction strengths represented in Fig. 8. Images were overlaid and figures were assembled using Adobe Photoshop (v. 20.0.6) and Illustrator CS (v. 10.14), respectively. Statistical analyses and plotting of data were conducted using RStudio (v. 1.1.463). Individual data points for each experiment were collected over multiple days. Statistical significance was determined using either a two-tailed Student’s t-test or Fisher’s exact probability test. Figure legends specify when each test was used and the p-value cut-off set.

## ACKNOWLEDGMENTS

We are grateful to R. Adikes, B. Kinney, A. Kohrman, M. Martinez and B. Martin for comments on the manuscript. We also thank I. Greenwald for providing the endogenously-tagged *lin-12*>LIN-12::GFP strain used in Fig. 7C.

## AUTHOR CONTRIBUTIONS

D.Q.M., T.N.M., and J.J.S. designed the experiments. All authors performed the experiments. T.N.M. and D.Q.M. performed the data analyses and wrote the paper with feedback from the other authors.

