## Supplemental Materials for "A developmental gene regulatory network for invasive differentiation of the *C. elegans* anchor cell"

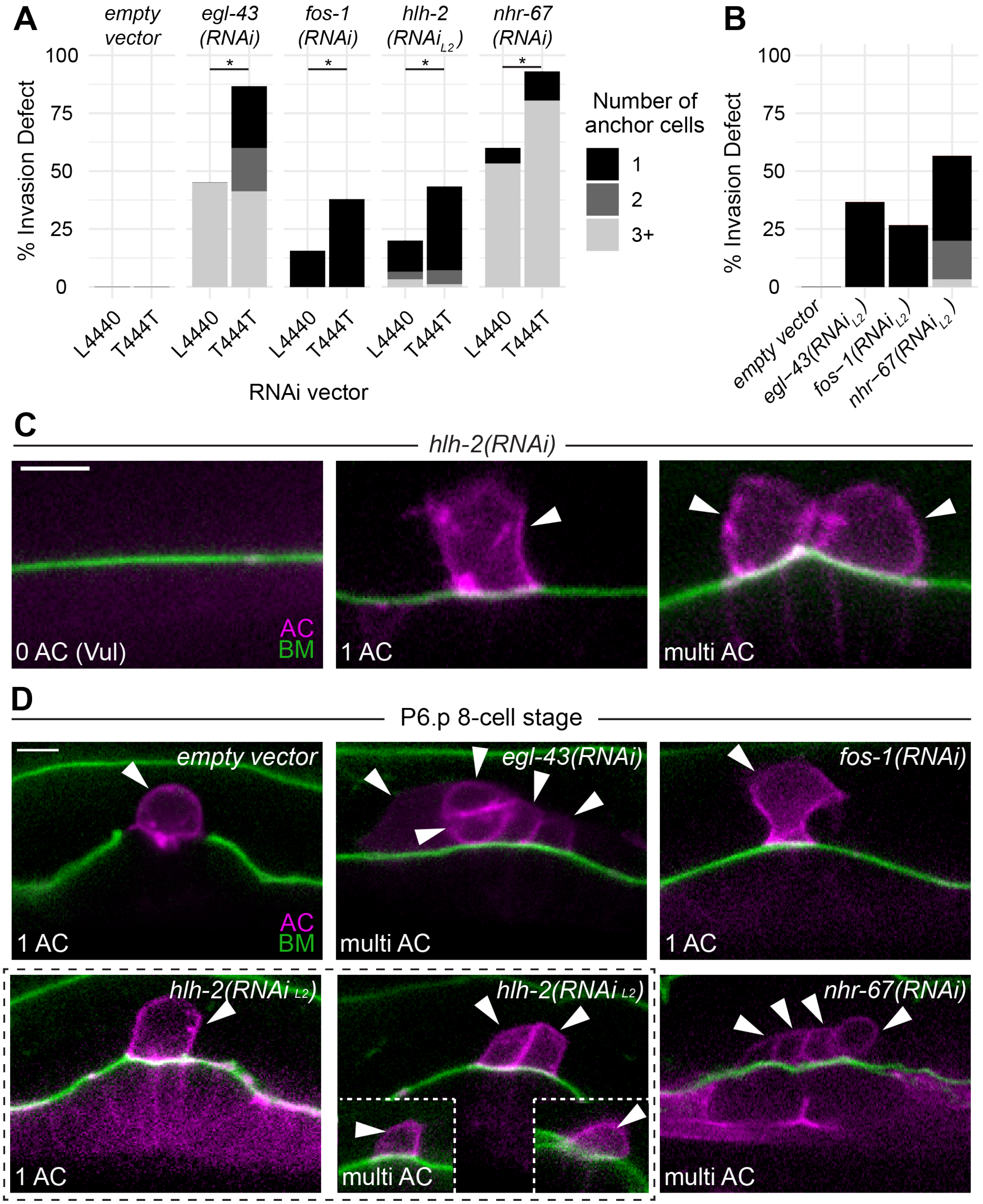
A
empty vector
egl-43 (RNAi)
fos-1 (RNAi)
hlh-2 (RNAi L2)
nhr-67 (RNAi)
% Invasion Defect
100
75
50
25
0
L4440
T444T
RNAi vector
Number of anchor cells
1
2
3+
B
empty vector
egl-43(RNAi L2)
fos-1(RNAi L2)
nhr-67(RNAi L2)
C
hlh-2(RNAi)
0 AC (Vul)
AC
BM
1 AC
multi AC
D
P6.p 8-cell stage
empty vector
egl-43(RNAi)
fos-1(RNAi)
hlh-2(RNAi L2)
nhr-67(RNAi)
1 AC
multi AC

**Fig. S1. Improved RNAi vector increases penetrance of TF depletion phenotypes.** (A) Stacked bar graph depicting the penetrance of AC invasion defects at the P6.p 4-cell stage comparing L4440-based versus T444T-based RNAi depletions in a uterine-specific RNAi-hypersensitive background. Asterisk (*) denotes a statistically significant difference between the vectors and represents a p-value < 0.03 by Fisher's exact test (n ≥ 30 animals per treatment). (B) Stacked bar graphs depicting AC invasion defects following delayed TF-RNAi treatment at the L2 stage (n = 30 animals per treatment). (C) Single plane of confocal z-stack depicting early depletion of *hlh-2* at the L1 stage. The AC-specific membrane marker (magenta, *cdh-3*$^{1.5}$>mCherry::PLCδ$^{PH}$) and BM marker (green, *lam-1*>LAM-1::GFP) are overlaid in each micrograph. (D) Single plane of confocal z-stacks during the L4 stage (P6.p 8-cell stage) depicting representative phenotypes associated with TF-RNAi treatment.

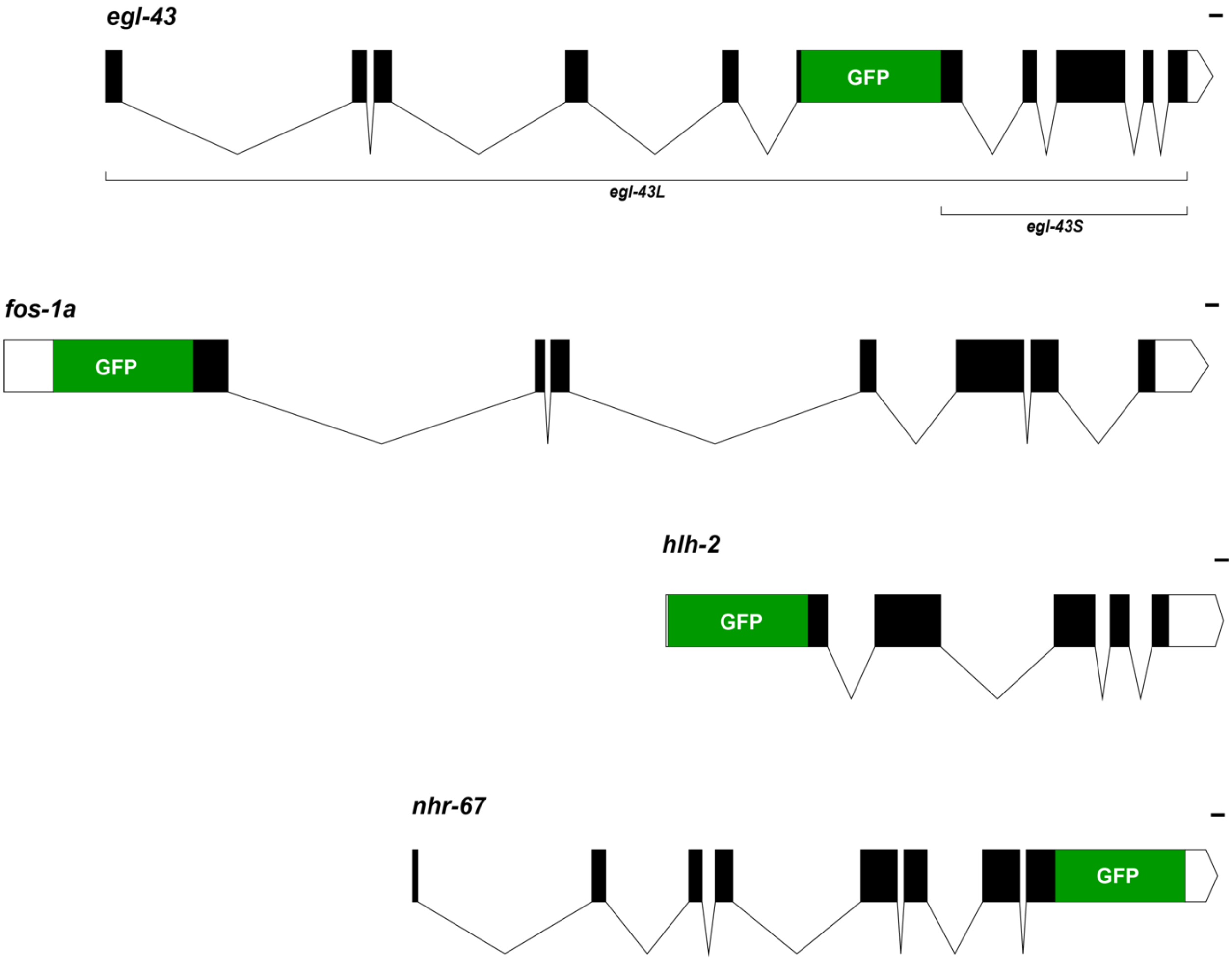


**Fig. S2. Schematic of endogenous GFP-tagged loci of pro-invasive TFs.** Codon-optimized GFP was integrated at the N-terminus of *fos-1A* and *hlh-2,* at the C-terminus of *nhr-67*, and internally in the *egl-43* locus (tagging both isoforms). This figure was made using http://wormweb.org/exonintron. Scale bar, 100 bp.

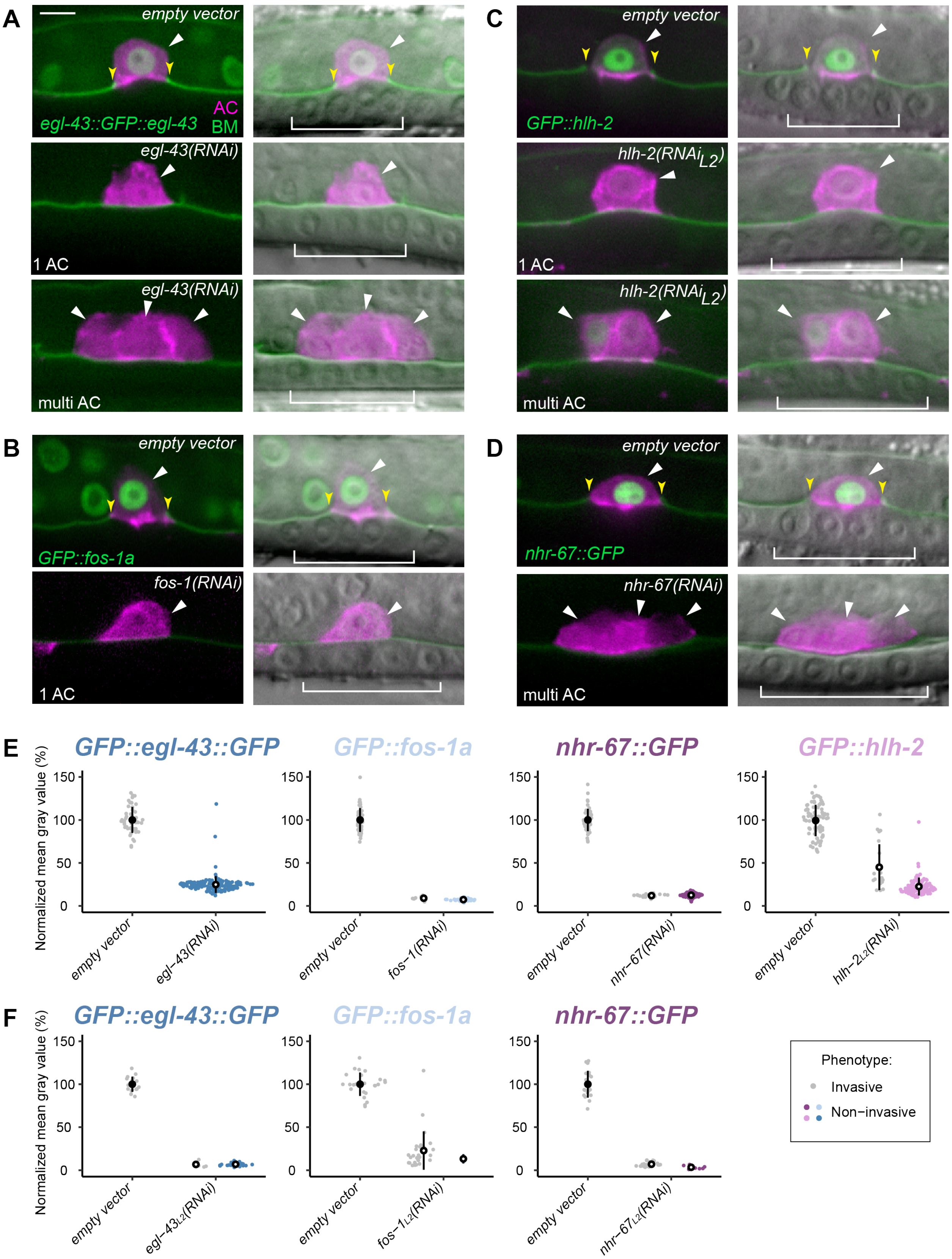
A
empty vector
egl-43::GFP::egl-43
AC
BM
egl-43(RNAi)
1 AC
egl-43(RNAi)
multi AC
B
empty vector
GFP::fos-1a
fos-1(RNAi)
1 AC
C
empty vector
GFP::hlh-2
hlh-2(RNAi L2)
1 AC
hlh-2(RNAi L2)
multi AC
D
empty vector
nhr-67::GFP
nhr-67(RNAi)
multi AC
E
GFP::egl-43::GFP
GFP::fos-1a
nhr-67::GFP
GFP::hlh-2
Normalized mean gray value (%)
150
100
50
0
empty vector
egl-43(RNAi)
fos-1(RNAi)
nhr-67(RNAi)
hlh-2L2(RNAi)
F
GFP::egl-43::GFP
GFP::fos-1a
nhr-67::GFP
egl-43L2(RNAi)
fos-1L2(RNAi)
nhr-67L2(RNAi)
Phenotype:
Invasive
Non-invasive

**Figure S3. AC invasion phenotypes do not necessarily correlate to TF expression levels.** (A-D) Single planes of confocal z-stack depicting representative phenotype (single vs. multi AC, bottom left of each image) of fluorescence alone (left; AC, magenta, expressing *cdh-3*>mCherry::moeABD and BM, green) and DIC overlay (right). (E) Sina plots of GFP-tagged TF levels, defined as the mean gray value of individual AC nuclei at the P6.p 4-cell stage following TF-RNAi knockdown. In this and all other figures, statistical significance as compared to empty vector controls is denoted as an open black circle and here represents a p-value of $< 1 \times 10^{-6}$ by Student's t test ($n \geq 50$ animals per treatment).

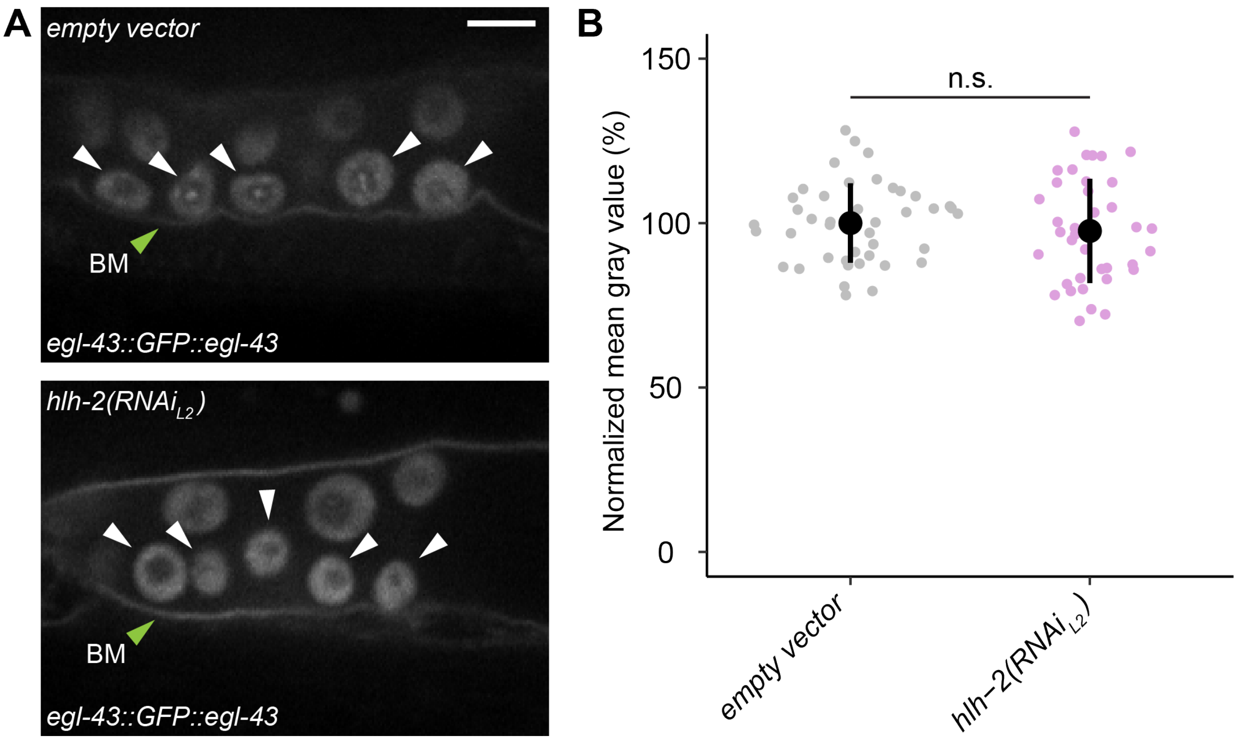


**Fig S4. HLH-2 does not regulate EGL-43 expression in the ventral uterus.** (A) Single plane of confocal z-stack depicting levels of *egl-43::GFP::egl-43* in the ventral uterine (VU) cells (white arrowheads) in empty vector controls (top) as compared to *hlh-2(RNAi)* depletion. BM indicated by green arrowheads. (B) Sina plots of *egl-43::GFP::egl-43* levels, defined as the mean gray value of individual VU nuclei, following *hlh-2* RNAi treatment (n ≥ 10 animals, n ≧ 37 VU cells quantified per treatment; n.s.*,* not significant, p-value = 0.45, Student's t-test).

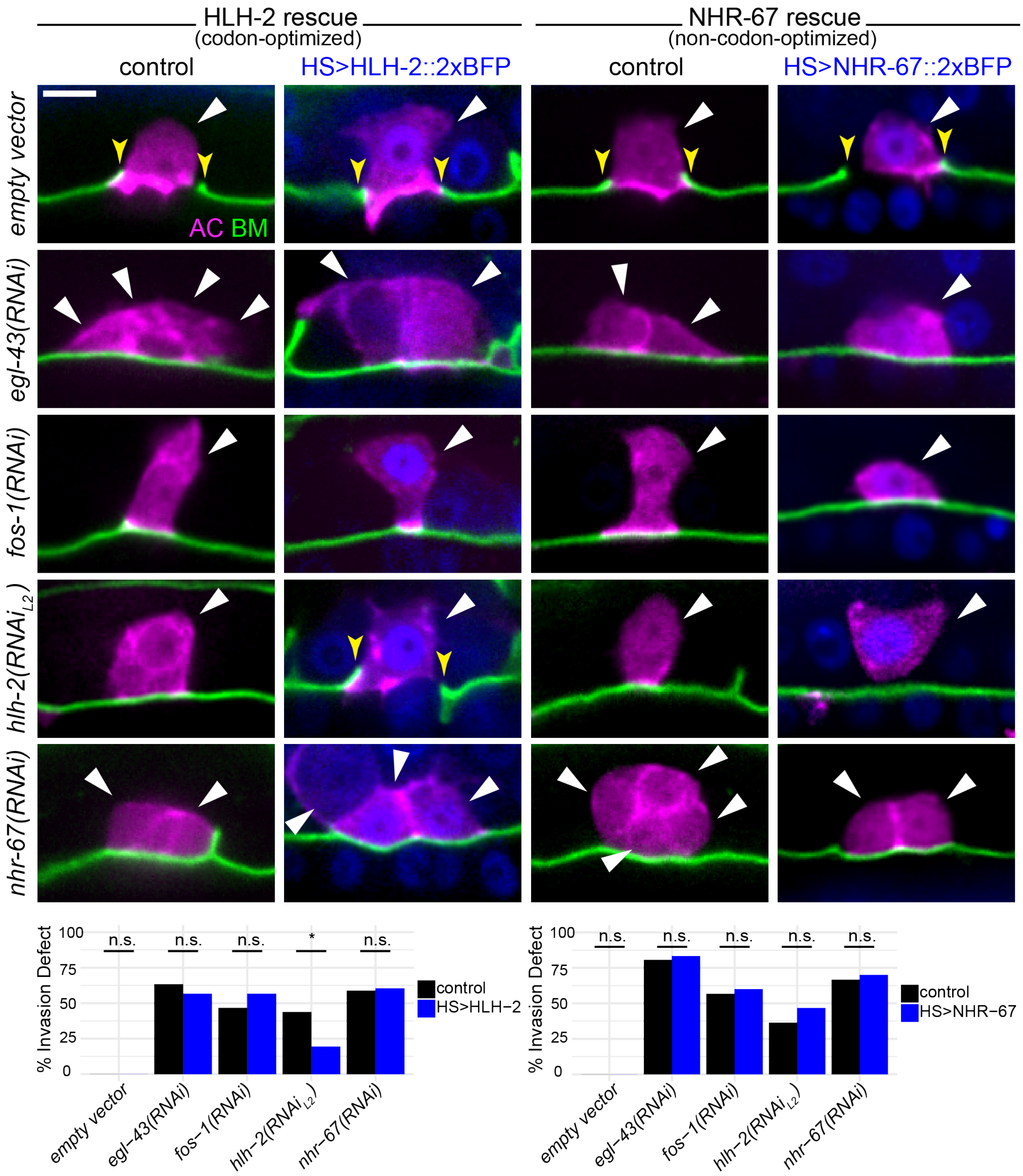


**Figure S5. Induced expression of *hlh-2* and *nhr-67* reveal cell cycle independent roles for *hlh-2*.** (A-B) Single planes of confocal z-stack depicting representative phenotype (single vs. multi AC) as fluorescence overlays (AC, magenta, expressing *cdh-3*>mCherry::moeABD, and BM, green) following no heat shock control (left) as compared to heat shock induced expression of HLH-2::2xTagBFP (blue) (A) and NHR-67::2xTagBFP (B). (C-D) Bar graphs depict the penetrance of invasion defects in control (black) compared to heat shock induced HLH-2 (C) and NHR-67 (D) at the P6.p 4-cell stage (n ≥ 30 animals examined for each RNAi treatment; n.s., not significant; *p-value < $1\times10^{-6}$; Fisher's exact test).

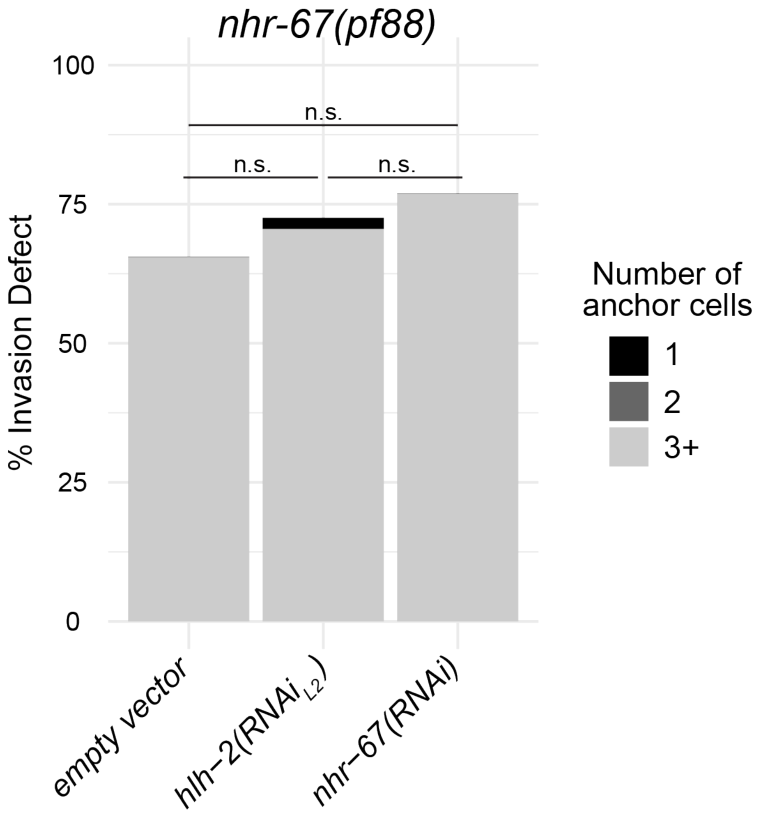


**Figure S6. Depletion of *hlh-2* does not significantly increase the invasion defect of an *nhr-67(pf88)* hypomorph.** Bar graph depicts the penetrance of AC invasion defects at the P6.p 4-cell stage in control (empty vector) as compared to *hlh-2(RNAi$_{L2}$)* or *nhr-67(RNAi)* treatment (n ≧ 50 animals examined for each RNAi treatment, n.s. not significant, Fisher's exact, p-value = 0.5350 (empty vector vs. *hlh-2*), 0.1690 (empty vector vs. *nhr-67*) and 0.6676 (*hlh-2* vs. *nhr-67*)).

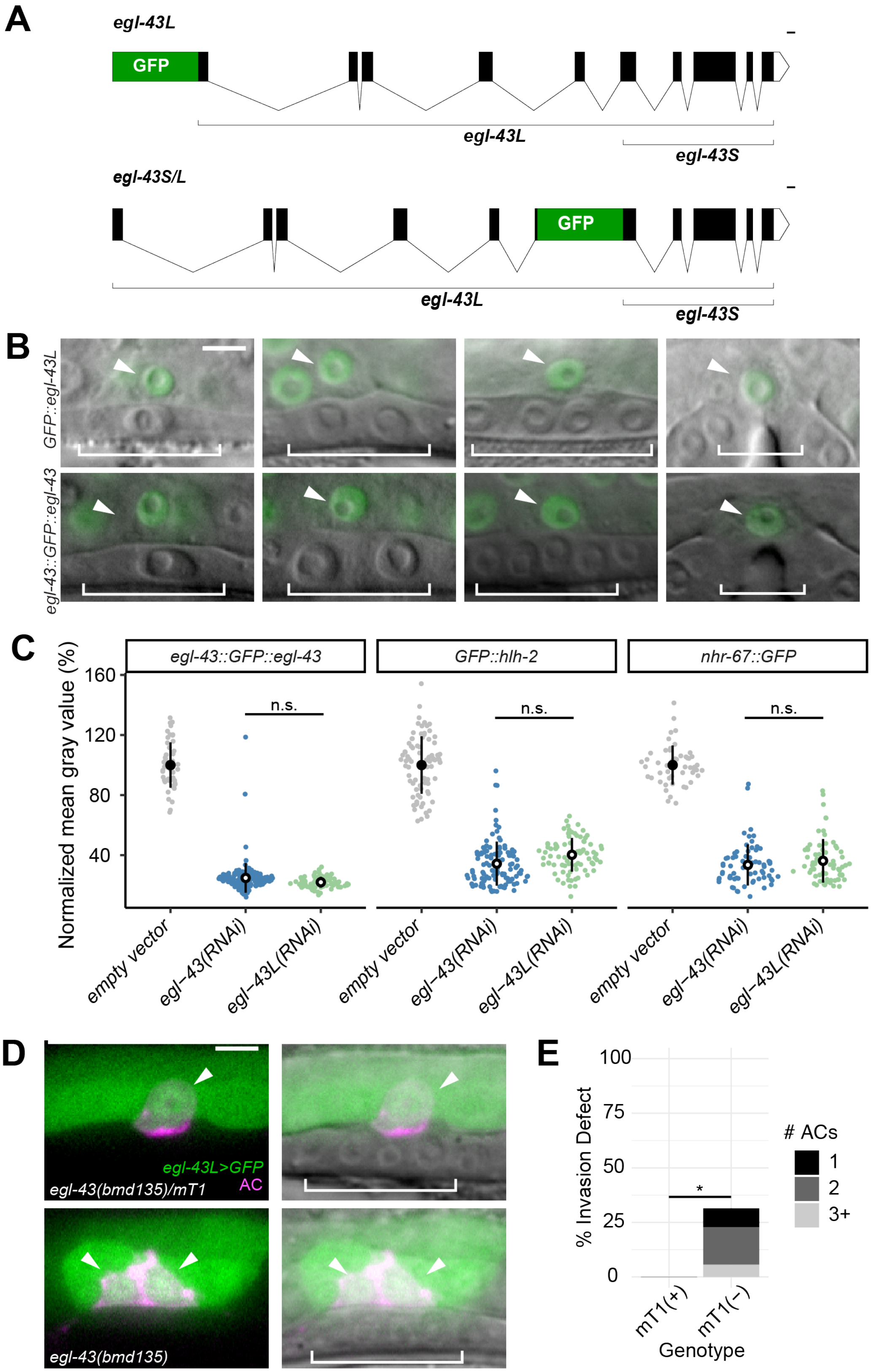
A
egl-43L
GFP
egl-43L
egl-43S
egl-43S/L
GFP
egl-43L
egl-43S
B
GFP::egl-43L
egl-43::GFP::egl-43
C
egl-43::GFP::egl-43
GFP::hlh-2
nhr-67::GFP
Normalized mean gray value (%)
160
120
80
40
n.s.
n.s.
n.s.
empty vector
egl-43(RNAi)
egl-43L(RNAi)
empty vector
egl-43(RNAi)
egl-43L(RNAi)
empty vector
egl-43(RNAi)
egl-43L(RNAi)
D
egl-43L>GFP
AC
egl-43(bmd135)/mT1
egl-43(bmd135)
E
% Invasion Defect
100
75
50
25
0
*
### ACs
1
2
3+
mT1(+)
mT1(−)
Genotype

**Figure S7. Both isoforms of *egl-43* function redundantly to regulate AC invasion.** (A) Schematics (via http://wormweb.org/exonintron) of GFP insertion into the *egl-43* locus to tag the long (top) or both isoforms (bottom). Scale bar, 100 bp. (B) DIC overlaid with single confocal planes of *GFP::egl-43L* (top) and *egl-43::GFP::egl-43* (bottom). (C) Sina plots of GFP-tagged TF levels, defined as the mean gray value of individual AC nuclei, following RNAi perturbation (n ≧ 25 animals for each treatment; n.s., not significant) between RNAi treatments targeting *egl-43L* and *egl-43*. (D) Single plane of confocal z-stacks depicting representative micrographs of control (top) versus *egl-43(bmd135)* animals expressing *egl-43L>GFP* (green) and an AC reporter (magenta). (E) Stacked bar graph depicting penetrance of AC invasion defect. Asterisk (*) denotes statistical significance between control and mT1(-) animals and represents a p-value $< 1\times10^{-3}$ by Fisher's exact test (n ≧ 25 animals per treatment).

#### Table S1: Scoring table

**percentages may not necessarily sum to 100 due to rounding*

| Genotype | RNAi Treatment (gene, vector, L1/L2 plating) | | | P6.p stage | % Invaded (#ACs) | | | | % Not invaded (#ACs) | | | | n |
|---|---|---|---|---|---|---|---|---|---|---|---|---|---|
| | | | | | 0 | 1 | 2 | 3+ | 0 | 1 | 2 | 3+ | |
| *rrf-3(pk1426) II; unc-119(ed3) III; rde-1(ne219) V; fos-1a>RDE-1, myo-2>YFP; cdh-3>PH::mCherry; lam-1>LAM-1::GFP + unc-119(+)* | *empty vector* | T444T | L1 | 4-cell | 0 | 100 | 0 | 0 | 0 | 0 | 0 | 0 | 50 |
| | *egl-43(RNAi)* | T444T | L1 | 4-cell | 0 | 12 | 1 | 0 | 0 | 27 | 19 | 41 | 75 |
| | *egl-43(RNAi)* | T444T | L2 | 4-cell | 0 | 63 | 0 | 0 | 0 | 37 | 0 | 0 | 30 |
| | *egl-43L(RNAi)* | T444T | L1 | 4-cell | 0 | 41 | 1 | 0 | 0 | 19 | 15 | 24 | 80 |
| | *fos-1(RNAi)* | T444T | L1 | 4-cell | 0 | 62 | 0 | 0 | 0 | 38 | 0 | 0 | 66 |
| | *fos-1(RNAi)* | T444T | L2 | 4-cell | 0 | 73 | 0 | 0 | 0 | 27 | 0 | 0 | 30 |
| | *hlh-2(RNAi)* | T444T | L1 | 4-cell | 0 | 30 | 0 | 0 | 21 | 37 | 6 | 5 | 115 |
| | *hlh-2(RNAi)* | T444T | L2 | 4-cell | 0 | 57 | 0 | 0 | 0 | 36 | 6 | 1 | 83 |
| | *nhr-67(RNAi)* | T444T | L1 | 4-cell | 0 | 7 | 0 | 0 | 0 | 13 | 0 | 81 | 72 |
| | *nhr-67(RNAi)* | T444T | L2 | 4-cell | 0 | 43 | 0 | 0 | 0 | 37 | 17 | 3 | 30 |
| | *empty vector* | T444T | L1 | 8-cell | 0 | 100 | 0 | 0 | 0 | 0 | 0 | 0 | 50 |
| | *egl-43(RNAi)* | T444T | L1 | 8-cell | 0 | 40 | 5 | 5 | 0 | 0 | 4 | 45 | 55 |
| | *egl-43L(RNAi)* | T444T | L1 | 8-cell | 0 | 78 | 9 | 5 | 0 | 7 | 0 | 2 | 58 |
| | *fos-1(RNAi)* | T444T | L1 | 8-cell | 0 | 90 | 0 | 0 | 0 | 10 | 0 | 0 | 50 |
| | *hlh-2(RNAi)* | T444T | L1 | 8-cell | 0 | 41 | 3 | 0 | 24 | 24 | 1 | 8 | 76 |
| | *hlh-2(RNAi)* | T444T | L2 | 8-cell | 0 | 64 | 1 | 1 | 0 | 23 | 1 | 9 | 81 |
| | *nhr-67(RNAi)* | T444T | L1 | 8-cell | 0 | 32 | 0 | 0 | 0 | 0 | 0 | 68 | 57 |
| | *empty vector* | L4440 | L1 | 4-cell | 0 | 100 | 0 | 0 | 0 | 0 | 0 | 0 | 30 |
| | *egl-43(RNAi)* | L4440 | L1 | 4-cell | 0 | 55 | 0 | 0 | 0 | 0 | 0 | 45 | 31 |
| | *fos-1(RNAi)* | L4440 | L1 | 4-cell | 0 | 84 | 0 | 0 | 0 | 16 | 0 | 0 | 32 |
| | *hlh-2(RNAi)* | L4440 | L1 | 4-cell | 0 | 69 | 0 | 0 | 9 | 16 | 0 | 6 | 32 |
| | *hlh-2(RNAi)* | L4440 | L2 | 4-cell | 0 | 80 | 0 | 0 | 0 | 13 | 3 | 3 | 30 |
| | *nhr-67(RNAi)* | L4440 | L1 | 4-cell | 0 | 40 | 0 | 0 | 0 | 7 | 0 | 53 | 30 |
| *cdh-3(5kb)>CKI-1::GFP; zmp-1>mCherry* | *empty vector* | T444T | L1 | 4-cell | 0 | 100 | 0 | 0 | 0 | 0 | 0 | 0 | 51 |
| | *egl-43(RNAi)* | T444T | L1 | 4-cell | 0 | 10 | 0 | 0 | 0 | 90 | 0 | 0 | 31 |
| | *fos-1(RNAi)* | T444T | L1 | 4-cell | 0 | 20 | 0 | 0 | 0 | 80 | 0 | 0 | 35 |
| | *hlh-2(RNAi)* | T444T | L2 | 4-cell | 0 | 92 | 0 | 0 | 0 | 8 | 0 | 0 | 77 |
| | *nhr-67(RNAi)* | T444T | L1 | 4-cell | 0 | 100 | 0 | 0 | 0 | 0 | 0 | 0 | 42 |
| *laminin::GFP; zmp-1>mCherry* | *empty vector* | T444T | L1 | 4-cell | 0 | 100 | 0 | 0 | 0 | 0 | 0 | 0 | 27 |
| | *egl-43(RNAi)* | T444T | L1 | 4-cell | 0 | 19 | 3 | 0 | 0 | 30 | 22 | 27 | 37 |
| | *fos-1(RNAi)* | T444T | L1 | 4-cell | 0 | 49 | 0 | 0 | 0 | 51 | 0 | 0 | 53 |
| | *hlh-2(RNAi)* | T444T | L2 | 4-cell | 0 | 56 | 0 | 0 | 0 | 31 | 3 | 10 | 59 |
| | *nhr-67(RNAi)* | T444T | L1 | 4-cell | 0 | 47 | 0 | 0 | 0 | 0 | 35 | 18 | 62 |
| *egl-43>LoxP::GFP::EGL-43; cdh-3>mCherry::moeABD; laminin::GFP* | *empty vector* | T444T | L1 | 4-cell | 0 | 100 | 0 | 0 | 0 | 0 | 0 | 0 | 68 |
| | *egl-43(RNAi)* | T444T | L1 | 4-cell | 0 | 0 | 0 | 0 | 0 | 8 | 14 | 78 | 158 |
| | *egl-43(RNAi)* | T444T | L2 | 4-cell | 0 | 16 | 0 | 0 | 0 | 78 | 0 | 6 | 32 |
| *hlh-2>LoxP::GFP::HLH-2; cdh-3>mCherry::moeABD; laminin::GFP* | *empty vector* | T444T | L1 | 4-cell | 0 | 100 | 0 | 0 | 0 | 0 | 0 | 0 | 83 |
| | *hlh-2(RNAi)* | T444T | L2 | 4-cell | 0 | 24 | 0 | 0 | 0 | 76 | 0 | 0 | 50 |
| *fos-1>LoxP::GFP::FOS-1; cdh-3>mCherry::moeABD; laminin::GFP* | *empty vector* | T444T | L1 | 4-cell | 0 | 100 | 0 | 0 | 0 | 0 | 0 | 0 | 74 |
| | *fos-1(RNAi)* | T444T | L1 | 4-cell | 0 | 18 | 0 | 0 | 0 | 9 | 5 | 68 | 120 |
| | *fos-1(RNAi)* | T444T | L2 | 4-cell | 0 | 93 | 0 | 0 | 0 | 7 | 0 | 0 | 30 |
| *nhr-67>NHR-67::GFP; cdh-3>mCherry::moeABD; laminin::GFP* | *empty vector* | T444T | L1 | 4-cell | 0 | 100 | 0 | 0 | 0 | 0 | 0 | 0 | 70 |
| | *nhr-67(RNAi)* | T444T | L1 | 4-cell | 0 | 15 | 1 | 0 | 0 | 0 | 11 | 73 | 130 |
| | *nhr-67(RNAi)* | T444T | L2 | 4-cell | 0 | 76 | 0 | 0 | 0 | 6 | 0 | 18 | 34 |
| *egl-43L>SEC::GFP::EGL-43; laminin::GFP; cdh-3>mCherry::moeABD;* | *N/A* | | | 4-cell | 0 | 67 | 3 | 0 | 0 | 8 | 17 | 6 | 51 |
| *egl-43L>SEC::GFP::EGL-43/mT1; laminin::GFP; cdh-3>mCherry::moeABD* | *N/A* | | | 4-cell | 0 | 100 | 0 | 0 | 0 | 0 | 0 | 0 | 25 |
| *nhr-67(pf88); qyIs227[cdh-3>mCherry::moeABD] I; qyIs7[laminin::GFP] X.* | *empty vector* | T444T | L1 | 4-cell | 0 | 34 | 0 | 0 | 0 | 0 | 0 | 66 | 58 |
| | *hlh-2(RNAi)* | T444T | L1 | 4-cell | 0 | 23 | 0 | 0 | 0 | 0 | 0 | 77 | 51 |
| | *nhr-67(RNAi)* | T444T | L1 | 4-cell | 0 | 27 | 0 | 0 | 0 | 1 | 0 | 71 | 65 |

**Table S2: Rescue experiments** **percentages may not necessarily sum to 100 due to rounding*

| Genotype | Condition | RNAi Treatment (gene, vector, L1/L2 plating) | | | P6.p stage | % Invaded | % Not Invaded | n |
|---|---|---|---|---|---|---|---|---|
| *bmd142[hsp>HLH-2::2xBFP] I; qyIs225[cdh-3>mCherry::moeABD] V; qyIs7[laminin::GFP] X.* | control | *empty vector* | T444T | L1 | 4-cell | 100 | 0 | 30 |
| | control | *egl-43(RNAi)* | T444T | L1 | 4-cell | 37 | 63 | 30 |
| | control | *fos-1(RNAi)* | T444T | L1 | 4-cell | 53 | 47 | 30 |
| | control | *hlh-2(RNAi)* | T444T | L2 | 4-cell | 56 | 44 | 32 |
| | control | *nhr-67(RNAi)* | T444T | L1 | 4-cell | 41 | 59 | 34 |
| | heat shocked | *empty vector* | T444T | L1 | 4-cell | 100 | 0 | 30 |
| | heat shocked | *egl-43(RNAi)* | T444T | L1 | 4-cell | 43 | 57 | 30 |
| | heat shocked | *fos-1(RNAi)* | T444T | L1 | 4-cell | 43 | 57 | 30 |
| | heat shocked | *hlh-2(RNAi)* | T444T | L2 | 4-cell | 81 | 19 | 31 |
| | heat shocked | *nhr-67(RNAi)* | T444T | L1 | 4-cell | 40 | 60 | 43 |
| *bmd121[LoxP::hsp>NHR-67::2xBFP] I; qyIs227[cdh-3>mCherry::moeABD] I; qyIs7[laminin::GFP] X.* | control | *empty vector* | T444T | L1 | 4-cell | 100 | 0 | 30 |
| | control | *egl-43(RNAi)* | T444T | L1 | 4-cell | 19 | 81 | 30 |
| | control | *fos-1(RNAi)* | T444T | L1 | 4-cell | 43 | 57 | 31 |
| | control | *hlh-2(RNAi)* | T444T | L2 | 4-cell | 64 | 36 | 33 |
| | control | *nhr-67(RNAi)* | T444T | L1 | 4-cell | 33 | 67 | 30 |
| | heat shocked | *empty vector* | T444T | L1 | 4-cell | 100 | 0 | 30 |
| | heat shocked | *egl-43(RNAi)* | T444T | L1 | 4-cell | 17 | 83 | 30 |
| | heat shocked | *fos-1(RNAi)* | T444T | L1 | 4-cell | 40 | 60 | 30 |
| | heat shocked | *hlh-2(RNAi)* | T444T | L2 | 4-cell | 53 | 47 | 30 |
| | heat shocked | *nhr-67(RNAi)* | T444T | L1 | 4-cell | 30 | 70 | 30 |

#### Table S3: Strains

| Strain | Genotype | Description | Figure(s) | Source |
|---|---|---|---|---|
| NK1316 | *rrf-3(pk1426) II; unc-119(ed3) III; rde-1(ne219) V; qyIs102[fos-1a>RDE-1, myo-2>YFP]; qyIs24[cdh-3>PH::mCherry]; qyIs10[lam-1>LAM-1::GFP with unc-119(with)]* | uterine-specific RNAi strain with AC and BM markers | 1, 6, S1 | Matus et al., 2015 |
| GS9129 | *arTi145[ckb-3>mCherry::H2B] II; lin-12(ar624[lin-12>LIN-12::GFP]) III.* | endogenous *lin-12* GFP reporter with somatic gonad marker | 2, 7 | Attner et al., 2019 |
| DQM337 | *egl-43(bmd88[egl-43>LoxP::GFP::EGL-43]) II; qyIs225[cdh-3>mCherry::moeABD] V.* | endogenous *egl-43* GFP reporter with AC marker | 2, S7 | This study |
| DQM497 | *fos-1(bmd138[fos-1>LoxP::GFP::FOS-1]) V.* | endogenous *fos-1* GFP reporter | 2 | This study |
| DQM352 | *hlh-2(bmd90[hlh-2>LoxP::GFP::HLH-2]) I; qyIs225[cdh-3>mCherry::moeABD] V.* | endogenous *hlh-2* GFP reporter with AC marker | 2 | This study |
| DQM291 | *nhr-67(syb509[nhr-67>NHR-67::GFP]) IV; qyIs227[cdh-3>mCherry::moeABD] I.* | endogenous *nhr-67* GFP reporter with AC marker | 2 | This study |
| DQM335 | *egl-43(bmd88[egl-43>LoxP::GFP::EGL-43]) II; qyIs225[cdh-3>mCherry::moeABD] V; qyIs7[laminin::GFP] X.* | endogenous *egl-43* GFP reporter with AC and BM markers | 2-3, 7, S3, S4, S7 | This study |
| DQM515 | *fos-1(bmd138[fos-1>LoxP::GFP::FOS-1]) V; qyIs227[cdh-3>mCherry::moeABD] I; qyIs7[laminin::GFP] X.* | endogenous *fos-1* GFP reporter with AC and BM markers | 2-3, 7, S3 | This study |
| DQM350 | *hlh-2(bmd90[hlh-2>LoxP::GFP::HLH-2]) I; qyIs225[cdh-3>mCherry::moeABD] V; qyIs7[laminin::GFP] X.* | endogenous *hlh-2* GFP reporter with AC and BM markers | 2-3, 7, S3, S7 | This study |
| DQM368 | *nhr-67(syb509[nhr-67>NHR-67::GFP]) IV; qyIs225[cdh-3>mCherry::moeABD] V; qyIs7[laminin::GFP] X.* | endogenous *nhr-67* GFP reporter with AC and BM markers | 2-3, 7, S3, S7 | This study |
| DQM7 | *qyIs330 [laminin::mCherry]; qyIs232 [CDT-1::GFP]* | G1 cell cycle phase reporter with BM marker | 4 | Matus et al., 2015 |
| DQM533 | *zmp-1(qy17[zmp-1::LoxP::GFP]) III; qyIs225[cdh-3>mCherry::moeABD] V* | endogenous *zmp-1* GFP reporter with AC marker | 4 | This study |
| DQM39 | *qyIs17 [zmp-1>mCherry] II; qyIs266[cdh-3(5kb)>CKI-1::GFP] V* | *zmp-1* mCherry reporter with AC-specific CKI-1 over-expression | 5 | Matus et al., 2015 |
| NK272 | *qyIs7[laminin::GFP]; qyIs17[zmp-1>mCherry]* | *zmp-1* mCherry reporter with BM marker | 5 | Matus et al., 2015 |
| DQM503 | *egl-43(bmd87[egl-43>SEC::GFP::EGL-43] II/mT1; qyIs227[cdh-3>mCherry::moeABD] I; qyIs7[laminin::GFP] X.* | endogenous *egl-43* transcriptional reporter (with SEC) balanced over mT1 with AC and BM markers | 6 | This study |
| DQM444 | *bmd121[LoxP::hsp>NHR-67::2xBFP] I; qyIs227[cdh-3>mCherry::moeABD] I; qyIs7[laminin::GFP] X.* | heat shock inducible NHR-67 with AC and BM markers | S5 | This study |
| DQM552 | *bmd142[hsp>HLH-2::2xBFP] I; qyIs225[cdh-3>mCherry::moeABD] V; qyIs7[laminin::GFP] X.* | heat shock inducible HLH-2 (codon-optimized) with AC and BM markers | S5 | This study |
| DQM266 | *nhr-67(pf88); qyIs227[cdh-3>mCherry:: moeABD] I; qyIs7[laminin::GFP] X.* | nhr-67 hypomorphic mutant with AC and BM markers | S6 | This study |
| DQM500 | *bmd135[egl-43L>SEC::GFP::EGL-43] II/mT1; qyIs227[cdh-3>mCherry::moeABD] I; qyIs7[laminin::GFP] X.* | endogenous *egl-43L* transcriptional reporter (with SEC) balanced with mT1 balancer with AC and BM markers | S7 | This study |
| DQM494 | *egl-43(bmd136[egl-43L>LoxP::GFP::EGL-43]) II.* | endogenous *egl-43L* GFP reporter | S7 | This study |
| DQM300 | *egl-43(bmd88[egl-43>LoxP::GFP::EGL-43]) II.* | endogenous egl-43 GFP reporter | Will be made available through the CGC (cgc.umn.edu) | |
| DQM497 | *fos-1(bmd138[fos-1>LoxP::GFP::FOS-1]) V.* | endogenous fos-1 GFP reporter | | |
| DQM311 | *hlh-2(bmd90[hlh-2>LoxP::GFP::HLH-2]) I.* | endogenous hlh-2 GFP reporter | | |
| PHX509 | *nhr-67(syb509[nhr-67>NHR-67::GFP]) IV.* | endogenous nhr-67 GFP reporter | | |

#### Table S4: Primers

| Primer | Primer sequence (5’→3’) | Primer type | Amplicon | Template |
|---|---|---|---|---|
| DQM657 | tcactatagggagaccggcaATG | Forward | *hlh-2* and *nhr-67* synthesized DNAs for T444T | Twist Biosciences gene fragments for *hlh-2, nhr-67* |
| DQM658 | attgggtaccgggcccc | Reverse | | |
| DQM688 | gagctcAGATCTatgagcatcgacacagacttc | Forward | BglII-*egl-43L*-XhoI for T444T | Twist Biosciences gene fragment for *egl-43L* |
| DQM689 | acgtacCTCGAGctgactttgacacgttgggc | Reverse | | |
| DQM720 | TCACTATAGGGAGACCGGCAATG | Forward | BglII-*egl-43*-SalI for T444T | *egl-43* IDT gBlock |
| DQM722 | GCCCCCCCTCGAGGTCGAACTTTTGGCACCGGAAC | Reverse | | |
| DQM708 | ggtttcgccacctctgacttg | Forward | colony PCR screening of T444T constructs | T444T-based constructs |
| DM191 | gtaatacgactcactatagggcgaattgg | Reverse | | |
| DQM433 | ACGTTGTAAAACGACGGCCAG | Forward | amplify left homology arm (universal) | Twist Biosciences gene fragments |
| DQM434 | CTCCAGTGAACAATTCTTCTCCTTTACTC | Reverse | amplify left homology arm (universal) | |
| DQM435 | GCGTGATTACAAGGATGACGATGAC | Forward | amplify right homology arm (universal) | |
| DQM436 | GAAACAGCTATGACCATGTTATCGATTTCC | Reverse | amplify right homology arm (universal) | |
| DQM751 | tcctattgcgagatgtcttGTCCACTCTCTTATATAGCAGGTTTTAGAGCTAGAAATAGC | Forward | *fos-1a* sgRNA | pDD122 |
| DQM747 | tcctattgcgagatgtcttGgatgctcatcctgaaaacttGTTTTAGAGCTAGAAATAGC | Forward | *egl-43L* sgRNA | pDD122 |
| DQM438 | tcctattgcgagatgtcttGaagtcagATGCCATCACAAGGTTTTAGAGCTAGAAATAGC | Forward | *egl-43* sgRNA | pDD122 |
| DQM440 | tcctattgcgagatgtcttGAGTTTTCAGAACCTCAATGGGTTTTAGAGCTAGAAATAGC | Forward | *hlh-2* sgRNA | pDD122 |
| DQM412 | AGATTGTACTGAGAGTGCACCATATGCGGTGTGAAATACCGCAC | Reverse | amplify sgRNA (universal) | pDD122 |

#### Table S5: Plasmids

| Plasmid | Base vector | Description |
|---|---|---|
| pTNM011 | pDD122 | *egl-43* internal sgRNA |
| pTNM012 | pDD282 | *egl-43::GFP::egl-43* repair template |
| pTNM013 | pDD122 | *fos-1* N-terminal sgRNA |
| pTNM014 | pDD282 | *GFP::fos-1* repair template |
| pTNM015 | pDD122 | *hlh-2* N-terminal sgRNA |
| pTNM016 | pDD282 | *GFP::hlh-2* repair template |
| pTNM046 | pDD122 | *egl-43L* N-terminal sgRNA |
| pTNM047 | pDD282 | *GFP::egl-43L* repair template |
| pTNM051 | pAP088 | heat shock inducible HLH-2 (codon-optimized) |
| pWZ172 | T444T | *egl-43* RNAi |
| pWZ173 | T444T | *egl-43L* RNAi |
| pWZ174 | T444T | *fos-1* RNAi |
| pWZ175 | T444T | *hlh-2* RNAi |
| pWZ176 | T444T | *nhr-67* RNAi |
| pWZ193 | pAP088 | heat shock inducible NHR-67 |

#### Table S6: CRISPR reagents

| Gene | Guide | Left homology arm sequence | Right homology arm sequence |
|---|---|---|---|
| *egl-43 (internal)* | aagtcagatgccatcacaag | CTAAGATATGAGAACCCGTTTACAGAGTCCTTTTA<br>TAGAATTTTGGTTTTTATAATAGAGATGTATGGAA<br>ACCGGGCAAAGTTAATTAGGATTCTTACAGCCAA<br>CAAGAAAATTTTAAGATAAGTAACGACCAAAACTT<br>GAGCGGAGTTGAAAGTTCAGTGCATTACAATTGA<br>AGTTTTTATTTATTATTTATTTATCTTGGGCTAGAA<br>TAGATGGAGCATCTTCACAGGACTGGACTTATAT<br>AATGGCTATCTGCCTGCCTGCCTACCTGCCTTTC<br>GTTTTATTTATACTGTATTTGCGCAATATAAACTCA<br>TGCATTTCTATTTGTTAGAAGTTAAAAAAAACAAT<br>ATTTAGAAAACGTTCACATGTACTTTGATAGTTGG<br>CTCGCATGTTGGCAGAAGGCAGGCACAGGGAAC<br>CCTGAAGGCACTTAGGCAGGTTCGCTGGACAAA<br>ATACCTGCTGATTGTGCTGATTTATTTTCACTAAT<br>ATAATCAGATATGAAACATGGACAATTGGGTACA<br>ATACTAATAGAGGGTGTACTGCTATATAACTTCTC<br>CGAATCAAAACTTAACAGTCATTCACATTCACTAT<br>CACGTGTTATCAATCAATTTTTTTCAGCGGTCTCA<br>AACAACACTCTCATATTCATTGCTCCTTAAAGCCG<br>TTTCGGTGTCATTTGTGTCCAAAGTCCTACACACA<br>ATTCTCAAATCTGTGCAGGCACCGGAGAGTTCAT<br>TCAGTGAGTTATTAATTTAGAGTTTCTAAGAATAA<br>ACTGAGCACTGTAATAATCTATCGTGCAATGCTC<br>CTGGGCTAAGAGTCCTTAAAGAGAGGATATAAAA<br>CTGAATAAACCCCATAATCTCACAACAGGGGGGA<br>AAACTCGAGGCGTGGTCAAGAAGGTTATTGTCCT<br>CTGTACTAATAAAAACGGAAACGATTTCAATGATG<br>TGCCCCCACCTGAGGTGGTCCCTTCCTTAATTGT<br>TTACCCCGTGAGAAAAGTGCAAGGAGCCATGATG<br>CTCCGCCCTCTTTCTTCTCAACCAATAAGACACG<br>CGCGCCGCCCATCCCTCGTGCATCTTCCCTTTGG<br>GGTGTGTCACAGGGCTCATTCTTGTGACGTGCG<br>CTTCCCACCTTTACCAGTGTGTTATCAATCTCCGT<br>CTTATTCATTAGTGAATAAATATTCCAGGACGGTT<br>GGACGTGCCCAACGTGTCAAAGTCAG | ATGCCATCACAAGCGGCACTAACGAAGCATCGAC<br>CAGTTTGCGAGATGACTGCACTCTACAAACCATTG<br>ATGGCGCAACTTGCCGGATTGAGCGGAGCAGGTG<br>GATTAGGTTCCGTGCCATATTGGCCACATATACTT<br>CAAATGGCAACACAGGTTAGTTATTATTTTGAAAAC<br>TCAACTATAGATACCGTCTCCCCTATTACTATTGCA<br>TCCCATTATTCAACTTTGACGCTTTATATTTCAGTT<br>TCCGTTTGTCCTACATGAAAATGATCAATTGGCCA<br>ATTTTTTATTATTTCTTTCTCTATAGTTTTGTAGTTG<br>ATAAAATAATCAAATCGTCATAATGAACCGAGATAT<br>AGCATTTCAAAGTGAAATAGTGGGCTGCAATAGAG<br>GAAATAAGGTAATTTTCCCTCACCCCCTTGACTATA<br>CAACACTCTTGAGAGTATCCATGGACAACAATCAT<br>CGTACCTTAATTAACCTACCTTAAGCCTCCTCCCA<br>ACCTCTCCATCTCTTCCGTTCAATATGATACCTTCA<br>TTTCCCGCATCTCGGGGGCTTAATCCTCTTTCCTT<br>CTCTTCCTCAAGCGGCTAATGAACATGCACTCATT<br>TTCAGGCACCACATTTCCCGCTTGCTTTTCTTGCC<br>GCCAATCCAGAAGCGTACAAACTGATGCAGCAGA<br>CGACGTGTGCATCTCCAGATGCCGAGTGTTCCAG<br>GTTTGTGAATGATTTATATGGTGAATCAAAAAAAAA<br>GTGGAGTAGTTCTCTTACTCTTATCAATAATTGATT<br>TATTCTATTCACTCTTTAGTACTAGTGAGAAGCTAA<br>TATATTGTATAGAAAACACAAAAAAAAATCAATTTA<br>TTCCAGTGGACATGCCAGTGAATCTTCACCAACTA<br>CAACTGAACCAGTCGACCTAACAGCTACACCAAAA<br>CCTCCTTCGACGTCTGAAATGGAAACTACATCAAA<br>GTCCGACGATGGAGAGGATCGTGACAGCATCGGA<br>GACTCTGGGAATGATGATGATGATGACTCAGAAGC<br>TGGAGTTCTAGATGAGTCGTCCACAACAACGTCAA<br>CGAAAAAGCGTCCGACATCTCACACAATTTCCGAC<br>ATACTCGCTGCTCCACAACTCGGTGCTCAGGCTTT<br>GAATTCGACGTTCCTTGGCATGCTTCAGCGCTCGC<br>TTAATTATAATCCAGCAGTTCCATCGCCTCACTCAT<br>TTC |
| *egl-43L (N-terminus)* | gatgctcatcctgaaaactt | TATTTTCCGTCCAGCAATTTACTTCCAAAAATTCA<br>AACTAAGTCAATAGTTGTTATAGTTTTTAAGTTGG<br>TCTTTGATTATTGAGCAGTTCGGTAAACTCGCAG<br>CCGACCTCGATGGAGTCTCGCGATAGCCTACTC<br>CTGGAAAAAACGTTTGTCTTCTGCTTTCACCTCAT<br>GATATCAGGAAACTTATTACAGACATTCAGCCGA<br>CAAATAAACGGCTTGGCTTCTACTAGAAATGCTC<br>AATAGATTCTAATTTAGTAGGTGTAGATATTTTCA<br>CTTTTGTTTATTTTTCTAAGTATCGAAGTACATATG<br>TTAGAAAGTAAATAAAAATTGCTATTCTACTTTAA<br>GAATTTTCTATATTAAAATAGTCTTTAGTAACGTG<br>GGGTTTTCGAAACTTGCAACTTTTCAAGTGAAGAT<br>GCTCCTCACCTGAAGCATCCTGTCACTCTACCGA<br>TTGGCTGATGGGCGTGTCGTCTTCCATCTCTTCC<br>ACCCGCCCACTTTTTGAGACAGGAGAGGCTCTC<br>GCTCCTCTCGCATCCACAAATGGAAGGCGCCTCT<br>TTCCATCTTCCTTTCTTTTTTTTTCTGGTTAGTTGAA<br>GTATCCTTAAAGTATCTTGACGTTTTTACTATATAT<br>GACTTCTGGAGAGACATTCGAGTTTTTGGATCAA<br>ATATAGAATGTCAAATTTTGAAGAGCAATCTTCTC<br>ATATGTCAATTTTTCAATAAAATTGTATCCTACATT<br>GTGGTGGTGTAGAACCCTCAAAAAATAATCTCTA<br>AATTAAGAATTTCTTGATTTTAAAACTGACTAGCA<br>AATTTGTCATGAAAAATCTCAAACGAATTTCCTAC<br>AGTTCATCTACCAAATGAATTGGGCTTGTAAACTT<br>TTTCACCATCTACTAAAAAATTCAATAAAATAATAA<br>TCTCTTCCCAAACTACACCTCCCTCCAATATCTCT<br>CAGCTTACTCCTAACTCCTATCCCAAATTCGTACA<br>TCTACCGTACCATTCCTCCTCCCTGTTCGTTCTCA<br>ATGACAAAGCCAGCCTTGAACAGCCATATCCAAC<br>TTTTTCGTTGCATTGTTCTCGTTCACAAGATGTAT<br>TACTCATCTTTCGCCCCAAAACTGTTCTTGTAATT<br>TGTACAGTTCTTGTTTCTTCCCACGTACAGTTTCC<br>TAAGTTTTCAGG | ATGAGCATCGACACAGACTTCCTCACGAGTGTTGA<br>GGTAAAGGAGGATGAGCTACATGGAAATGTGCTC<br>ATTGCAGTAACTCAAATTGCACTCGGAAGGACCAT<br>CGGTGTGATTGATAAGGTGAGCTTTTGGTGTACAA<br>TTTTTACGCCGTAGACATTGTCTTAGAGGTGTCCG<br>CTAGAAAAAGAAGTATAAGCTTATTCCCGGGCGTC<br>TAAAAAACTCCACCGGAAAAAATACAGGTCGCTTG<br>AACTGAAATATCTTATAGCTTGAGTAACCTATGAAA<br>TAAGCTTTCAGATGTTACACTCATATTTTGTTAGTT<br>AGTAGTTATATGATAACAGTAATTTTAGGTTGTAAT<br>TGATAAAATAATGACTTTTTTAAATTAACAAAACTTT<br>TTTTAAAATATTTTTACAAGCATTTTAAATTGATACC<br>TTGGTGCCTAATCTACCTAATCTACAGTACCTTCG<br>CCAACATTTTCAAATCCTTTCCTAATGATTCAACGT<br>TCGAATTACAAATTCTTCCCAGAACCCAAAACCGC<br>AATTATCGGGACCCTGAAATGTTTGGCGCCATGTG<br>TTCCTCCCTGCTTCTTCTCCTCAAAAATTCCTTACA<br>CCTGTCTTTCCACCTTTCCCGCTCCCCCCGTCCAT<br>TACCACTTTGTTAACCTTGGGGGCACACTAGTAAC<br>CATTACATTAGGATTTA |

| | | | |
|---|---|---|---|
| *fos-1A (N-terminus)* | ccactctcttatatagcaga | CTTAGACATCCATCCAAATATTTAAATGTTGCTGT<br>TGCTTTTGGGTCAACAAACGACACATACCTGTTA<br>GTGTCATCCTTCGTTCCGCGTGCACGCATCGCAC<br>TGAATCATCGGTATTCATCACCTCTTCTCTGCAAT<br>CAACGAGCCAAATGCCAATTGGGTTATTTGAGTT<br>AGAAACTCGGGAAGCTTCACAGAAGAAATAATTA<br>GCTGAATTGGCAAAGAGAGCACCTGCTCCCCGG<br>AATGTCATACTTTTGTAGAATAAAAGAAGGGAAGA<br>CGACGACGTCTCCGGTTCTTCTATCCCAAACAAT<br>ACCATTTTTTGTTTCCCATCTCGTCTTCGATGTTG<br>CTGACTCACCATCAATATCTAAACAGAAAGCACC<br>CAATGGTTCGCCTTTTTTGTCTCCTTTCTCGACAG<br>TTTCTGTTTTTTTTTTGTATGTTACATAAATACCGTA<br>CACCCACCATCTAGTCAATTGTCACATATTCGAAA<br>TCAAAAGTTATCAAAAACTCTCGTGCACTTTAATA<br>CACAGTTCCGTCGGCCACCCCGTCACCAGCCGG<br>CGTTTTCTTGGTGATATCATTGGCGGTGGCTCTT<br>TGAAAATTCAAATTTGGTGTCGTCCTCCCGAAGG<br>TGTGGGCGGAGCTTGCGCACCGTTCGCGGTGGC<br>CGTTGCTCTCGACAGTTTTCTCTTTCCCTCGCTC<br>CTTTCTTCTCTCAACCAAACACCTGCCCACACAC<br>GCAACTTACTTTTGTCATCGCTGGTCCCCTGTCC<br>AAAGATACAATCAACCCGTGTTGTGCTAATCATCA<br>CCTAGAAGAGTCATCTTCGTGAATCAGAATCCCT<br>TTTCTTCCCCCTACTTTTTCCACTTCTATTCCCGG<br>TTACTCACCTTCTGTGTTTGGCCCCGATCGTGTG<br>TCAGCACGTCTGCTATATAAGAGAGTGCA | ATGTTCGAACAACCATCGTCGACGACTAACACCAC<br>CACGAGCTCTGGTTCTGGCTCCGATTCCAATCACT<br>ACTTCGAATTGGGTCCCAGGAACCCGATCAACCAA<br>GCGCACCCGACATCAGTCATTGTTCCACCGCGAC<br>AACATCACCATCAGATCCACCAACAACAAACCGAC<br>AACTCACCTCTAACTCCGTGTACACCATATTATCCA<br>TCAAATGCATATGGACTCCCTTTATTCTTTGGAACT<br>GATTTCCTGCAGGTATGATACATCACAAATATCTTC<br>CTTAATTCAGTGTTTACTTCAAAGTCCACTATTCCC<br>TTGCTAATAATCACGAGTTTTGATAGAGCTTTTAAA<br>TCATTACTTTCACTCATTTTTCACTCAACCTCGAGC<br>CTTACATTTGTCAGTTGATCTCTTCTTTCCCATATA<br>AAAATGAGATTATAGTAGCTTACTTTCATTTAGTTT<br>AATATTAATTCTGAAAGTATCAAACCAAAAAGTAAA<br>GCGTTCACTAATTCTTTTGTGCAAAAGTCATGCAG<br>AAAAAAGTTACCTAGCTCTTGAATCTTTTCCCTCAA<br>AAATGTTCTTACCAGCCATCTCCCCCGGATTTTAG<br>TTCCGGAAAAATAAAACTATTTGGTTATGGATTAAA<br>TGTATATGGATAGGTGGTGCACGAGGAATCTTAAA<br>AGAAGATGCCGACGTGTTTACACACTTCTACAGTA<br>ATCTATCTATCTTTCACCTTTCCCCCAATTGTTACT<br>TTTGTGATCTTCTTTACCTCTCATGCACTCGCTAAT<br>CCGTTTTGGATTGTTTCCTGTTTGCACCTGTTGAAA<br>AAAGATAAGTAATACTTATATGTTAACTTTTTTCGG<br>TGCCAGTGAAAATATTACAGGCACGCAAATTTTTTA<br>AATATTCAAAAAATATCTGTCACTTCTGCTTTGAGC<br>TTG |
| *hlh-2 (N-terminus)* | agttttcagaacctcaatgg | GACTGGTGAATGAATGGTGGGAGAAGAGATGGA<br>GGACCCCGGGATAATAGAGGATTGTGTGCTGCA<br>GGAGGTAGAGAGAGATGGTTGTGGGCGTGACCT<br>CTCCCGATCATCGTCTCTGCGTCTCTTGGATGTA<br>GACGGTCCCCGGGAGAAAGAATGTCTGCGTCTA<br>TGTACACCAATATACTGTTTATGTCCCTTGAATGG<br>GCTTTTATACGCGTCCCTCCACTACCACCACCCT<br>CGAAGAACCATTCCTAGCCACCCGCATAGCCGCT<br>AAGAAGGGGGGAGTGCACTCTTTTCGTCTTTTTT<br>TGTGTCTAAATCTGTTCCGTTTACTAGCTTTTCAA<br>GTTCATGATCTTACTGTAATAAAACATAGCACATA<br>TTTTGAATGAAATACATAATGTATCCCATCTTCTG<br>ATTTTGAAAATTAAAATTTCTCTCCTCCCAATTTTC<br>AATTTTGGATCTTTTACCAAATAATTTTGATCTGCA<br>TTACTTTTTCTCACTCCTACAATGTTTACGAATCC<br>TTACCAATTTTGAATTTAAACAAGAACGCAAATGT<br>ATTGTAGGGCAGTTTTTTTTTCAATTATTGAGTTTA<br>TCAAAAAATGTATATTTGCATCCGAAATGTTACTA<br>GCTCACGTTTAAAGCTTTCATATATATCTTATCCT<br>ACGTCTGAATTTAAACTAATTATTTAGATGGGCCT<br>GAAACCCTTTGATCCATCACCCCTTTACTGTTATT<br>TTATTGCTCTTTCGGGATGTCATCCATCCGATAAT<br>CATCTACAATGACATCTACATTTACAACTATTTCTT<br>GATACCATCTCAACATCATTACCTCCCTACAAATT<br>GTTCTCCTTCAAGTAACTTGCTCCTAATAATTTAT<br>ACCGGAACTTAGTAGATCTATTGTGACGTCACAA<br>TTATTATTCTTTATTGATCTACCTTGCCTCTTTTGA<br>TCTCGTTTGCCTATTTAACCAAACGAGGAGAATC<br>GATTATTTTCTTCTTTTTCTTATTCTCAATATTTCTT<br>CAAATGGGGTTGTCTTGACGAAAATCTAATCAGA<br>ATATATTTACTAACCTAATTTTTTCACCTGCTGCTC<br>CAGCCCAATTCCCTACTTATTCCCTTTCCTCTCCC<br>CGCTTCTCCGCTGACTTTATTTCTTTGACATTACT<br>ACTATAATCGTCTTTCATATCATTTCGCAGAAATA<br>AAGTTTTCAGAACCTCA | ATGGCGGATCCAAATAGCCAACTTACGTCAGCCAC<br>AACTGTTGCAACTGCCGCCATTGCTCAACCACAGG<br>TTATGCTTCCAAATGCATATGATTATCCTTATAATA<br>TTGATCCGACAACGATTCAGATGCCTGATTATTGG<br>AGTGGTTAGTCTTTTTTTCATTTGAAATCTATTATAA<br>TTTCAATTATATCATTTGGTAAACCAAACCAAACAT<br>GTTTTCGTGAATTTGTTAAAAGTCGCTTGTAATCAG<br>TTAGAAATATTGGATAAATTGCAGAAAAAGGATATA<br>AGTTTGGTATTACTTATCTCGAATTGTTGAATTGTT<br>GATTTTATAAGCTCGCAAATTTTTAAAATTTACTTTA<br>AGTTGAAACAAAATGTATAACTCTTTTAATGTTCTA<br>AAATTTTGTAAAAATCTGGTTTGCTCTAGTAAGTAG<br>TTTATTAACCAAAATGTTTATATCTCATTTCTCAAGT<br>GCAAGTCATCTAAATAAACATTTTCAGGATACCATC<br>TCAACCCGTATCCTCCGATGCAAACAACGGATATT<br>GATTATTCATCAGCCTTCCTTCCAACACATCCACC<br>AACTGAAACCCCTGCTTCCGTAGCTGCTCCAACTT<br>CTGCAACATCTGATATTAAGCCAATTCATGCAACAT<br>CATCCACTTCAACGACGGCTCCATCTACTGCTCCA<br>GCTCCAACTTCAACTACTGATGTGCTTGAGTTAAA<br>GCCAACAACAGCTCCAGCCACGAATTCTGCAGAA<br>ACATCAGCGATTGTTGCTCCACAGCCTCTTACTAA<br>TCTTACCGCACCAATTGACGCAATGTCATCAATGT<br>ATACATGGCCACAAACATATCCAGGTTACCTTCCA<br>CCTTCAGAAGATAACAAAGCAAGTGAAGCTGTTAA<br>TCCATACATCTCAATCCCTCCAACATATACATTTGG<br>TGCTGATCCATCAGTTGCCGACTTCTCATCGTATC<br>AGCAGCAACTTGCTGGACAGCCGGTACGTTTCTAT<br>TCTTCTTGGTTTTATTCGTGTTTTACTGTTTTACCA<br>CAAGTTAATCTCGACGAATGATTGCTTGGCCCTCC<br>TCAGCCGCCAAGCATTCACGTATGCATAATCCGCT<br>TCCTTTTGATGTGAGCCGCCGCCTTCTGAAGATTG<br>AGATGCATGCAAAAGGCGGCATCTTCTTGAGAAGA<br>ACACCCTGTGCGCACTAAACGACCTCC |
| *nhr-67 (C-terminus)* | ccacgtcattcgattcgatcaat<br>agagagtgttaatgttgaagagg | N/A (made by SunyBiotech) | N/A (made by SunyBiotech) |

#### Table S7: RNAi vectors

| RNA Target | Synthetic gene block |
|---|---|
| *egl-43L(RNAi)* | TCACTATAGGGAGACCGGCAATGAGCATCGACACAGACTTCCTCACGAGTGTTGAGGTAAAGGAGGATGAGCTACATGGAA<br>ATGTGCTCATTGCAGTAACTCAAATTGCACTCGGAAGGACCATCGGTGTGATTGATAAGGCTACACCGAATGATTCGAATGC<br>TCTTCTCATCTTGAACCTGATTAAGGAAGCTGATGACGGAGAAGATGCCAATATTTGTATGAGGCAGGAGGATAGAAAGACT<br>TTTCTACAAACCAGTAAGATTATCAATATTGGAGAGCGTCTTCTTCTACAAAGACTGTCCGAAGAGGAGTGTGATGAGGAGG<br>ATCAGGATGATCTTGAGAATTTAATTTTGTTAAAAGATGAAGATCGGCCGGACAGTACTCAAAGCTGCACAAAGAGCAGCAG<br>TGAAGACAGCAATCTAAACGGTTTCGAGGAGTATATTCGAGAACACGGCGAACTTGTACCTGGTCAAACGCCTCCCGACGG<br>ATCACACAAGTGTGGAGTTTGTCCAAAGAGTTTTTCAAGTGCAAGCGGTCTCAAACAACACTCTCATATTCATTGCTCCTTAA<br>AGCCGTTTCGGTGTCATTTGTGTCCAAAGTCCTACACACAATTCTCAAATCTGTGCAGGCACCGGAGAGTTCATTCAGACGG<br>TTGGACGTGCCCAACGTGTCAAAGTCAGGGGGGGGCCCGGTACCCAAT |
| *egl-43(RNAi)* | TCACTATAGGGAGACCGGCAATGCCATCACAAGCGGCACTAACGAAGCATCGACCAGTTTGCGAGATGACTGCACTCTACA<br>AACCATTGATGGCGCAACTTGCCGGATTGAGCGGAGCAGGTGGATTAGGTTCCGTGCCATATTGGCCACATATACTTCAAAT<br>GGCAACACAGGCACCACATTTCCCGCTTGCTTTTCTTGCCGCCAATCCAGAAGCGTACAAACTGATGCAGCAGACGACGTG<br>TGCATCTCCAGATGCCGAGTGTTCCAGTGGACATGCCAGTGAATCTTCACCAACTACAACTGAACCAGTCGACCTAACAGCT<br>ACACCAAAACCTCCTTCGACGTCTGAAATGGAAACTACATCAAAGTCCGACGATGGAGAGGATCGTGACAGCATCGGAGAC<br>TCTGGGAATGATGATGATGATGACTCAGAAGCTGGAGTTCTAGATGAGTCGTCCACAACAACGTCAACGAAAAAGCGTCCG<br>ACATCTCACACAATTTCCGACATACTCGCTGCTCCACAACTCGGTGCTCAGGCTTTGAATTCGACGTTCCTTGGCATGCTTC<br>AGCGCTCGCTTAATTATAATCCAGCAGTTCCATCGCCTCACTCATTTCTAAGAGCAATGAGCGGAGCAAAAGCATCATCATC<br>ACCAAGCTCGAGCAGTGGAAGTGGAAAGGATAGGTACACGTGCAAGTTCTGTCAGAAAGTGTTCCCAAGATCAGCGAATTT<br>GACAAGACATTTAAGGACACATACAGGAGAGCAGCCTTATAAGTGTCAATATTGCGAGCGAAGCTTCTCAATCTCCTCCAAT<br>TTGCAACGCCACGTTAGAAACATTCATAATAAGCCGAACACCTCGCTGACACCCCACAATCATCATCGTCAACGAAGTCTGC<br>ACAATTCAACCTCAACCTCCACTACTACTACTACCGTCCATCATCCTCTTCTTCATCTTCCAGGAACGTCCGTTCCGGTGCCA<br>AAAGTTACCGGGCCCCCCCTCGAGG |
| *fos-1(RNAi)* | CTCACTATAGGGAGACCGGCAGATCTATGGTACAGTATAGCACGGTGAAGAAATCGTCGGCTGGGAGAAAACCTAAAGAAG<br>AGGATAATATGGAGGATGACGATGATGATAAGAGGCTGAAACGTCGTCAAAGGAATAAAGAAGCTGCTGCAAGATGTCGGC<br>AAAGGAGAATTGATTTGATGAAGGAATTGCAAGATCAAGTAAATGACTTCAAAAATAGCAACGACAAAAAAATGGCGGAATG<br>CAACAACATCCGAAATAAGCTCAACAGTCTCAAAAACTATTTGGAAACGCATGATTGTAAACTGAGTCGCGAAGAACGAACA<br>CATGAGATAAATCGATTGATAATCCCACCATCAACAGTTCCACCATCACAACCATATCTTCAACATTCGTTGAGAGTTCATCC<br>ACCACGTGCTGACTCTGTACCATATTCTATTCGATCAGGACATTCATCTTCATCGTCTGAACAACACTCACCAGTTGAGGACT<br>ATAAGCCTTCAATTGATCAATTACTTCTTCCTCCAATCTCATGTATTCAAAATATCAAAGATCGAAATATCAATTCTATGCCACC<br>ACCAGCTTTACCGGCAAGCACGAGTGCTGCTGGAATTCATGTTATCACTTCTATCCCGGTTTCGCATGCCAACTCATTACAC<br>GGTCGTTCGGAAAACGTTTTCGCAGAACCGGAACGCAAAATTCCAAAAATCGAGTTGGACCAAACACTGACATCATTGACCA<br>TGCCAGACGACGTTGAACGTCCGTCAGCTCTTCCGACGCTTTCCAGAATCGTTGAAAATCAGCCGATCACTACTCCCAGCA<br>GGCCTTTCCGTCTCGGGGGGGAATATCAAAATCAAACACCGCAGAGTACAGGGAATGGACTATTCGGAGGCCCACCAGGC<br>CCATTCGACTTGCTTTCCTCCAACACTGGTCTGACACCTTCTGGTCAGCCAACAATGAACTTCGTTTCAACTCCAACCCCGAT<br>TCAACCACATCCGGATGCTGATCTCCGACCACTCTAGGTCGACCTCGAGGGGGGGCCCG |
| *hlh-2(RNAi)* | TCACTATAGGGAGACCGGCAATGGCGGATCCAAATAGCCAACTTACGTCAGCCACAACTGTTGCAACTGCCGCCATTGCTC<br>AACCACAGGTTATGCTTCCAAATGCATATGATTATCCTTATAATATTGATCCGACAACGATTCAGATGCCTGATTATTGGAGT<br>GGATACCATCTCAACCCGTATCCTCCGATGCAAACAACGGATATTGATTATTCATCAGCCTTCCTTCCAACACATCCACCAAC<br>TGAAACCCCTGCTTCCGTAGCTGCTCCAACTTCTGCAACATCTGATATTAAGCCAATTCATGCAACATCATCCACTTCAACGA<br>CGGCTCCATCTACTGCTCCAGCTCCAACTTCAACTACTGATGTGCTTGAGTTAAAGCCAACAACAGCTCCAGCCACGAATTC<br>TGCAGAAACATCAGCGATTGTTGCTCCACAGCCTCTTACTAATCTTACCGCACCAATTGACGCAATGTCATCAATGTATACAT<br>GGCCACAAACATATCCAGGTTACCTTCCACCTTCAGAAGATAACAAAGCAAGTGAAGCTGTTAATCCATACATCTCAATCCCT<br>CCAACATATACATTTGGTGCTGATCCATCAGTTGCCGACTTCTCATCGTATCAGCAGCAACTTGCTGGACAGCCGAATGGTC<br>TTGGTGGAGATACCAACTTGGTTGACTACAATCATCAATTCCCACCAGCCGGTATGAGCCCACACTTTGATCCAAATGGATA<br>TCCAGGAATGACCGGAATGCCACCAGGATCAAGTGCATCATCTGTTCGAAATGATAAGTCTGCATCAAGAGCAACGAGCCG<br>TCGTCGGGTACAAGGCCCTCCATCTTCTGGAATTCCAACTCGCCATTCATCGTCTTCAAGGCTTTCTGATAATGAATCAATGA<br>GTGATGACAAAGATACGGATAGGAGATCACAGAATAATGCTCGAGAAAGAGTACGAGTTCGTGACATCAGGGGGGGCCCG<br>GTACCCAAT |
| *nhr-67(RNAi)* | TCACTATAGGGAGACCGGCAATGATGACTGCGGTTTCACAGATGTCGGTTCCAAGCAGTCGAATTCTATTGGATGTTGATTG<br>TAGAGTCTGTGAAGATCATTCGTCGGGGAAGCATTACAGTATTTTTTCCTGCGACGGGTGTGCTGGATTCTTTAAACGTTCAA<br>TCCGTCGTCATCGTCAATACGTGTGCAAAAACAAAGGCAGTCCATCCGAAGGACAATGCAAAGTGGACAAAACACACCGCA<br>ACCAGTGCAGAGCTTGCCGACTAAGAAAGTGTCTGGAAATTGGGATGAACAAAGATGCTGTCCAACATGAGAGAGGGCCTC<br>GTAACTCAAGTTTAAGACGACAACAGATGATGTTCGATCATGGATCCTCTCCTAATTCGCCGGAAATGGGATCAGAAAGTGA<br>TGCAATAATTCTTCCGACATCATCAATGAATCGTGACACAGTGGCCGGTACTGCTGCTCGAATCTTTTTTGCACTTGTTGGAT<br>TTTGTCAGAATCCTTTGAATGGAGTACCTAAAGAACGGCAAATGACAATGTTTCAACAAAATTGGGCAGCACTTCTTGTTCTC<br>CATGCAACTGAAACACGTGCCATAACTTCTAAGCAAATAAGGACTGAGACGATATCCGGTAGCTCAGAGCAAAGAAATGCG<br>GTTGCTAACGCGTTTGAAATAATTGAACGTTTACAATTGGATAATAGAGAATACATGATGTTGAAACATTTCACAATGTGGAG<br>AGATACACCAAGTGCAATTCAAATTGTCTTTCAGTTAGCATCTATTCAGAATTTCACACATAGAACCGAACCGACGAGATATA<br>TCCAATGTATAAATGCAATTGCCGCTATTCCAACAACTTCAATAATTGATGTTCTATTCCGCCCTTCAATTGGATCAGCTTCAA<br>TGCCAAGACTTATTCAAGACATGTTCAAGCCACCACAACAACCCACTCCTACGTCACTGTTTCCAATGGGGGGGGGCCCGG<br>TACCCAAT |
